# MitoArc: A Reproducible Short-Read Workflow for Human Mitochondrial Genome Analysis with Circular-Coordinate and NUMT-Aware Evaluation

**DOI:** 10.64898/2026.11.19.767817

**Authors:** Sarvesh Galgale, Isha Vats, Agnik Haldar

**Author notes:** **Corresponding author** Agnik Haldar. **Equal contribution** Sarvesh Galgale and Isha Vats contributed equally to this work.

## Abstract

Human mitochondrial DNA (mtDNA) analysis from short-read sequencing is complicated by nuclear mitochondrial DNA segments (NUMTs), circular-reference coordinates, sequencing depth, and variant-caller configuration. We developed MitoArc, a Nextflow workflow that accepts paired-end FASTQ files or pre-aligned BAM and CRAM files, retains available nuclear mapping context, and performs standard and shifted-coordinate processing. It runs Genome Analysis Toolkit (GATK) Mutect2 and mutserve2 independently and reports mitochondrial quality metrics, NUMT evidence, haplogroups, contamination estimates, functional annotations, consensus sequences, and reproducibility outputs. We evaluated MitoArc using 18 controlled datasets, spanning three technical replicates and six sequencing depths (100×–5000×), with 648 allele-level truth observations. Requiring confident primary mitochondrial mapping eliminated 33 observable synthetic NUMT-associated false-positive calls but reduced recall from 0.894 to 0.796. In a matched evaluation comprising 108 truth observations, standard-only and standard-plus-shifted GATK processing produced identical variant recovery and allele-fraction estimates. GATK recall increased from 0.741 at 100× to 0.917 at 5000×, while temporary working storage increased approximately 20-fold. Against the complete truth set, GATK recovered 548 observations and mutserve2 recovered 311 under the evaluated configurations; mutserve2 insertion and deletion modes were not enabled. The benchmark used synthetic data without a background sequencing-error model. A reduced HG002 dataset demonstrated end-to-end operation of the pre-aligned workflow, subject to incomplete nuclear context. MitoArc preserves caller-specific evidence and the analytical context needed to interpret mitochondrial variants. The software is publicly available at https://github.com/sarvezh1/mitoarc.

## 1. Introduction

The human mitochondrial genome is only 16,569 bp, but its analysis from short-read sequencing data is not straightforward. Mitochondrial DNA (mtDNA) is present in multiple copies within a cell, and different alleles can coexist within the same individual as heteroplasmy. The fraction of molecules carrying a variant can differ between tissues and over time, making accurate estimation of mitochondrial allele fractions important in studies of inherited disease, aging, population variation, and somatic mutation (Stewart and Chinnery 2021). Deep sequencing has made low-level mitochondrial variation increasingly accessible, but it has also placed greater demands on read mapping, variant calling, and the interpretation of low-frequency signals.

Several properties of the mitochondrial genome create specific problems for short-read analysis. First, the mitochondrial genome is circular, whereas standard sequence aligners and variant callers operate on linear reference sequences. The artificial breakpoint introduced by linearization can therefore affect reads spanning the ends of the reference sequence. Second, the nuclear genome contains numerous sequences of mitochondrial origin, commonly referred to as nuclear mitochondrial DNA segments or NUMTs. Sequence similarity between mtDNA and NUMTs can cause mitochondrial reads to align to nuclear loci or nuclear-derived reads to align to the mitochondrial genome, producing false-negative, false-positive, or biased heteroplasmy estimates (Maude et al. 2019). Large-scale analysis of 56,434 genomes in gnomAD further showed that NUMT-derived false-positive calls are influenced by mitochondrial copy number and can form a substantial fraction of apparent low-level heteroplasmies. That analysis used both standard and shifted mitochondrial references together with additional filtering intended to limit NUMT-associated artifacts (Laricchia et al. 2022). More recent long-read pangenome analysis has also shown that human NUMTs include substantial polymorphic and non-reference variation that is incompletely recovered by conventional short-read approaches, indicating that nuclear-mitochondrial sequence ambiguity cannot be represented fully by a single reference genome (Fu et al. 2026).

These problems have motivated several dedicated human mtDNA analysis tools. MToolBox accepts raw or pre-aligned sequencing data, reconstructs mitochondrial genomes, estimates heteroplasmy, assigns haplogroups, and annotates detected variants; its mapping procedure also realigns mitochondrial reads against the nuclear genome to remove reads attributed to NumtS (Calabrese et al. 2014). mtDNA-Server introduced scalable heteroplasmy analysis together with quality control, annotation, and contamination assessment, and its later mtDNA-Server 2 implementation added a Nextflow workflow, expanded input validation, coverage assessment, and a calling mode combining mutserve2 SNVs with Mutect2 indels (Weissensteiner et al. 2016, 2024). MitoHPC was developed for whole-genome sequencing data and estimates mtDNA copy number and heteroplasmy while addressing circular-reference coverage, recovering reads from selected NUMT regions, and generating a sample-specific mitochondrial consensus for a second round of variant analysis (Battle et al. 2022). More recently, MitoGEx combined quality control, GATK mitochondrial variant calling, annotation, haplogroup classification, phylogenetic analysis, and visualization within a graphical workflow for human mitochondrial sequencing data (Jeenkeawpiam et al. 2026). Integrated mtDNA analysis pipelines therefore already exist, and the value of another workflow cannot rest on integration alone.

A separate question is how individual analytical choices affect the final result when they are evaluated under the same conditions. NUMT filtering may remove nuclear-derived signals but can also remove genuine mitochondrial reads. Shifted-reference processing addresses a known property of the circular mitochondrial genome, but its incremental effect cannot be isolated when the standard-only and shifted analyses differ in calling interval, filtering, or other post-processing. Increasing mitochondrial depth can improve the recovery and quantification of low-frequency alleles while also increasing computational demand. Variant callers may also differ in sensitivity, allele-fraction estimation, supported variant classes, and default configuration. When these choices occur together within different pipelines, their individual effects can be difficult to separate. This makes controlled, matched evaluation useful in addition to conventional end-to-end software validation.

Here, we present MitoArc, a Nextflow DSL2 workflow for reproducible analysis of human short-read mitochondrial sequencing data from paired-end FASTQ, BAM, or CRAM input. MitoArc performs full-reference alignment for FASTQ input and preserves the available mapping context of pre-aligned BAM/CRAM input before mitochondrial read extraction, applies standard and shifted-coordinate mitochondrial processing, and runs GATK Mutect2 and mutserve2 as separate callers rather than combining their results into a single consensus callset. The workflow also produces mitochondrial coverage measurements, experimental NUMT evidence, haplogroup and contamination estimates, functional annotation, caller-specific consensus sequences, static figures, MultiQC output, and a self-contained HTML report.

Evaluation of MitoArc focused on four methodological questions. First, we asked whether requiring confident mitochondrial mapping before variant calling reduces observable NUMT-associated false-positive signals and whether this affects recovery of true mitochondrial variants. Second, we tested whether standard-plus-shifted processing provides additional boundary-specific recovery when downstream calling and filtering are matched between processing modes. Third, we examined how variant recovery, allele-fraction error, and computational requirements change across mitochondrial sequencing depths from 100× to 5000×. Fourth, we compared GATK Mutect2 and mutserve2 against the same controlled truth panel under explicitly fixed caller configurations. These experiments used three independently generated technical replicates at each of six sequencing depths, together with a separate HG002 real-data analysis to evaluate operation of the complete downstream workflow. The aim was not to identify a universally optimal caller, filtering strategy, or sequencing depth, but to make the consequences and limitations of these analytical choices explicit and measurable.

## 2. Materials and Methods

### 2.1 Workflow design and implementation

MitoArc v1.0.0 was implemented as a Nextflow DSL2 workflow for short-read analysis of the human mitochondrial genome. Nextflow was used to define process dependencies, manage intermediate files, and support reproducible execution across computational environments (Di Tommaso et al. 2017). The release was validated with Nextflow 26.04.6. Software dependencies were assigned to version-pinned Docker containers. The evaluated MitoArc v1.0.0 release is archived on Zenodo (https://doi.org/10.5281/zenodo.23154158).

The workflow accepts paired-end FASTQ, coordinate-sorted BAM, or CRAM input. FASTQ data undergo quality control, read preprocessing, full-reference alignment, and sorting before entering the common downstream analysis. BAM and CRAM files enter through a separate validation step and are converted to the same internal alignment structure. Subsequent mtDNA extraction, coverage analysis, circular-coordinate processing, variant calling, interpretation, and reporting are shared across the three input routes.

GATK Mutect2 and mutserve2 are run as independent mitochondrial variant callers. Their calls remain separate throughout the workflow and are compared descriptively rather than merged into a single callset. MitoArc also produces mitochondrial coverage metrics, optional NUMT evidence, haplogroup and contamination estimates, functional annotations, caller-specific consensus sequences, plots, MultiQC output, and a self-contained HTML report.

### 2.2 Input handling and reference requirements

Each MitoArc run accepts one samplesheet format: paired-end FASTQ (sample,fastq_1,fastq_2), BAM (sample,bam), or CRAM (sample,cram). Mixed input types within a single samplesheet arerejected. Sample identifiers must be unique, and paired FASTQ entries must refer to separate read files.

FASTQ analysis is intended for alignment against a human reference FASTA containing the specified mitochondrial contig, which is chrM by default. The reference is indexed for BWA-MEM2 and samtools within the workflow working directory.

Pre-aligned BAM and CRAM files are checked before downstream analysis. Validation includes samtools quickcheck, coordinate sort order, presence of the requested mitochondrial contig, and agreement between the lengths of mapped alignment contigs and the supplied reference FASTA. CRAM files must also be decodable using the supplied reference. Missing alignment indexes are generated within the workflow working directory. BAM files are retained in BAM format, whereas CRAM files are decoded to BAM before entering the common downstream analysis.

Read-group sample identifiers are also inspected. A missing SM value or a value differing from the samplesheet identifier is recorded as a warning, whereas files containing more than one distinct biological SM value are rejected.

The alignment is then classified according to its retained nuclear context. Files containing all 22 canonical autosomes are labeled FULL_REFERENCE; those containing a subset are labeled PARTIAL_NUCLEAR_CONTEXT; and alignments without canonical autosomal sequence are labeled MT_ONLY. Analyses that require complete nuclear context are not inferred when the necessary information is absent.

### 2.3 FASTQ preprocessing and full-reference alignment

Raw paired-end FASTQ files are first assessed with FastQC 0.12.1. Read preprocessing is performed with fastp 0.23.4 using its default filtering and adapter-processing settings; MitoArc specifies the paired input and output files and requests the corresponding HTML and JSON reports (Chen et al. 2018).

Preprocessed reads are aligned to the supplied human reference using BWA-MEM2 2.2.1 (Vasimuddin et al. 2019). Alignment is performed using the number of CPUs assigned by Nextflow. A read group is added during alignment with the sample identifier used for both the read-group ID and sample field and with the sequencing platform recorded as ILLUMINA. The resulting SAM file is coordinate sorted and indexed using samtools 1.20 (Li et al. 2009).

FASTQ reads are aligned against the full human reference before mitochondrial reads are selected. This retains the initial competition between mitochondrial and homologous nuclear sequence during mapping and preserves nuclear alignment information required for later context-dependent analyses. Nuclear mitochondrial DNA segments can interfere with mtDNA read mapping and heteroplasmy detection, particularly for low-frequency calls, making the retained nuclear context relevant to subsequent NUMT assessment (Laricchia et al. 2022).

### 2.4 mtDNA extraction and coverage metrics

For each standardized alignment, MitoArc first identifies read names with at least one alignment to the mitochondrial contig. All alignment records associated with those read names are retained in a sample-specific mtDNA BAM and indexed with samtools. This preserves mate information associated with mitochondrial alignments before mitochondrial-only realignment.

Mitochondrial coverage is calculated from the extracted BAM using mosdepth 0.3.14 (Pedersen and Quinlan 2018). A minimum mapping quality of 20 is applied, and coverage is restricted to the configured mitochondrial contig. Per-base depth is used to calculate mean, median, minimum, and maximum mtDNA depth and the proportion of mitochondrial bases covered at ≥1×, ≥10×, ≥100×, ≥500×, and ≥1000×.

Nuclear coverage is estimated separately from the standardized input alignment. In the absence of a user-specified autosomal list, canonical autosomes (1–22 or chr1–chr22, according to the reference naming convention) are selected. Mean autosomal depth and a length-weighted median of chromosome-level depth estimates are calculated with the same mapping-quality threshold. If no autosomal sequence is available, or if the alignment retains a complete autosomal header but mapped reads are present for only a subset of those autosomes, the autosomal estimate is reported as unavailable.

When a valid mean autosomal depth is available, MitoArc reports a coverage-derived mtDNA copy-number proxy as

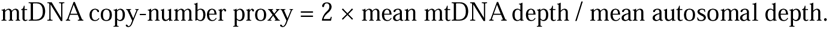

The value is reported as a sequencing-derived proxy rather than a direct measurement of mitochondrial copy number per cell.

### 2.5 Circular-coordinate processing

The human mitochondrial genome is circular, whereas sequence alignment and variant-calling tools operate on a linear reference representation. This creates an artificial boundary at the ends of the linearized mitochondrial sequence and can affect reads that span that boundary. Shifted mitochondrial references have therefore been used to move the artificial breakpoint during variant analysis (Laricchia et al. 2022).

For circular-coordinate processing, MitoArc converts the extracted mtDNA BAM back to paired FASTQ after read-name collation. Singleton and unpaired outputs are not retained. The mitochondrial sequence is then extracted from the supplied reference and used to construct two references: the original mitochondrial sequence and a shifted representation. The default shift offset is 8,000 bp. For a sequence of length *L*, the shifted reference consists of positions 8,001–*L* followed by positions 1–8,000. The shift offset must be greater than zero and smaller than the mitochondrial sequence length.

Both representations are indexed and the mitochondrial read pairs are independently realigned with BWA-MEM2 2.2.1. The resulting alignments are coordinate sorted and indexed. Read-group information contains the sample identifier and the ILLUMINA platform designation.

MitoArc assigns complementary calling intervals to the two representations. The standard-reference branch covers positions shift_offset + 1 through *L*, while the shifted-reference branch covers the final shift_offset bases of the shifted sequence. After coordinate conversion, the shifted interval corresponds to positions 1 through shift_offset in the original coordinate system. With the default 8,000-bp offset and a 16,569-bp mitochondrial reference, canonical positions 1–8,000 are therefore obtained from the shifted branch and positions 8,001–16,569 from the standard branch. Each canonical position is represented by one calling branch.

The use of an 8,000-bp shift is consistent with the shifted-reference approach used in the gnomAD mitochondrial workflow to address the artificial breakpoint of the linearized mitochondrial genome (Laricchia et al. 2022).

### 2.6 GATK Mutect2 mitochondrial variant calling

Mitochondrial variants are called with GATK 4.6.2.0 Mutect2 using --mitochondria-mode. Mutect2 is run independently on the standard and shifted mitochondrial realignments, with each invocation restricted to its corresponding complementary interval. Mitochondrial-mode Mutect2 has been used for high-depth human mtDNA variant analysis and for the detection of both homoplasmic and heteroplasmic variants (Laricchia et al. 2022).

Variants generated from the shifted alignment are converted to the original mitochondrial coordinate system using GATK LiftoverVcf and a rotation-specific chain file generated by MitoArc. A liftover containing rejected variant records is treated as an error rather than silently excluding those records. Mutect2 statistics from the standard and shifted branches are combined with MergeMutectStats, and the two non-overlapping VCFs are joined with GatherVcfs to produce a single standard-coordinate raw VCF. The combined VCF is indexed and validated against the standard mitochondrial reference.

Final GATK filtering is performed once, after coordinate conversion and combination, using FilterMutectCalls with --mitochondria-mode. MitoArc v1.0.0 does not supply an autosomal-coverage value to FilterMutectCalls and does not add an explicit minimum allele-fraction argument to the Mutect2 or filtering commands. NUMT-associated evidence is assessed separately by the experimental strategies described in Section 2.8 rather than through GATK’s autosomal-coverage filter.

The resulting raw and filtered GATK VCFs are retained separately. GATK calls are not merged with mutserve2 calls to generate a hybrid callset.

### 2.7 mutserve2 variant calling

mutserve2 2.0.3 is used as an independent mitochondrial variant caller. The caller is run on the standard mitochondrial realignment and its matching standard mitochondrial reference. A separate shifted-reference mutserve2 branch is not used. mutserve2 is the mitochondrial SNV caller used within mtDNA-Server 2 and is designed for the detection of homoplasmic and heteroplasmic mitochondrial variants (Weissensteiner et al. 2024).

The minimum heteroplasmy level is set to 0.01 through the mutserve2 --level option. The mitochondrial contig name and number of processing threads are also supplied explicitly. Other native calling settings are left unchanged in the v1.0.0 configuration, including a mapping-quality threshold of 20, base-quality threshold of 20, alignment-quality threshold of 30, and disabled base alignment quality (BAQ) adjustment. Native raw evidence output is retained together with the native VCF and call table.

The evaluated MitoArc v1.0.0 configuration does not enable mutserve2’s beta insertion or deletion options. The mutserve2 results reported in this study therefore apply to this fixed configuration and should not be interpreted as a general assessment of all mutserve2 indel-capable modes. This distinction is retained in the benchmark interpretation because mtDNA-Server 2 also distinguishes mutserve2-based SNV calling from Mutect2-based indel calling in its combined analysis mode (Weissensteiner et al. 2024).

MitoArc applies no additional biological filtering to the native mutserve2 calls. For downstream comparison, the VCF is normalized using BCFtools 1.20 (Danecek et al. 2021). The sample identifier is standardized, multiallelic records are split, reference alleles are checked against the mitochondrial reference, and records are coordinate sorted, compressed, and indexed. The native files are retained unchanged alongside this normalized representation. Because the mutserve2 VCF provides total depth and allele frequency but not an exact allele-depth field, MitoArc records alternate-read depth as unavailable rather than estimating it from allele frequency and total depth.

### 2.8 NUMT evidence strategies

Nuclear mitochondrial DNA segments (NUMTs) can complicate short-read mtDNA analysis because reads originating from homologous mitochondrial and nuclear regions may not map unambiguously. This can affect both variant detection and estimated heteroplasmy (Laricchia et al. 2022; Maude et al. 2019). MitoArc therefore includes an optional experimental NUMT framework whose interpretation depends on the retained nuclear context. FULL_REFERENCE alignments provide the intended whole-reference mapping context, whereas NUMT evidence generated from PARTIAL_NUCLEAR_CONTEXT inputs is treated as context-limited and is not used to infer complete-WGS NUMT performance. Three strategies are evaluated. The default mapping-quality threshold for this analysis is 20.

Strategy A represents the baseline caller output. No additional mapping-context filter is applied before mitochondrial read preparation and variant calling. GATK therefore follows the standard-plus-shifted calling path described above, while mutserve2 is run on the standard mitochondrial realignment. These results provide the reference callsets against which the other NUMT strategies are compared.

Strategy B applies a mapping-confidence filter before repeating the mitochondrial calling paths. The retained input alignment is examined by read name. A read name is retained when at least one primary mitochondrial alignment has a mapping quality of at least 20; secondary and supplementary alignments do not satisfy this retention criterion. All alignment records associated with retained read names are written to a new BAM. Mitochondrial read preparation, realignment, GATK calling, and mutserve2 calling are then repeated from this filtered alignment. An audit table records each read-name decision and the highest qualifying primary mitochondrial mapping quality. Strategy B therefore tests the effect of requiring confident primary mitochondrial mapping before variant calling; it does not require the fragment to have no nuclear alignment whatsoever.

Strategy C does not generate a new callset. Instead, it classifies the retained mapping evidence associated with reads supporting individual mitochondrial alleles. For each SNV, MitoArc identifies primary mitochondrial alignments carrying the alternate base and links their read names back to the retained input alignment. Supporting fragments are classified as HIGH, AMBIGUOUS, or LOW NUMT evidence. HIGH evidence is assigned when a fragment has a primary nuclear alignment at or above the mapping-quality threshold, or a qualifying secondary or supplementary nuclear alignment. Secondary alignments recorded through XA tags are treated as ambiguous nuclear evidence because these tags do not provide mapping quality, whereas supplementary alignments recorded through SA tags contribute HIGH evidence when their reported mapping quality meets the threshold. Lower-confidence nuclear alignments, a mate mapped to a nuclear contig, absence of a qualifying primary mitochondrial alignment, or failure to recover the read in the retained input alignment result in an AMBIGUOUS classification. LOW evidence is assigned when a qualifying primary mitochondrial alignment is present without these nuclear mapping signals.

At the variant level, a called SNV is marked FLAGGED_NUMT_EVIDENCE when at least two supporting fragments are observed and none is classified as LOW; other called SNVs are marked RETAINED. The fragment-support procedure is restricted to SNVs in v1.0.0, so indels and other non-SNV alleles are returned as UNRESOLVED_SUPPORT. Strategy C does not remove variants from the caller VCFs. Its status is added to downstream annotation as supporting evidence and is not treated as proof that a variant is false. Further details of Strategy C evidence classification and unresolved non-SNV support are provided in Online Resource 1 (Section S3).

### 2.9 Haplogroup and contamination assessment

Haplogroup classification is performed independently for the filtered GATK callset and the normalized mutserve2 callset using HaploGrep 3.3.2 (Schönherr et al. 2023). HaploGrep assigns mitochondrial haplogroups from observed variants using a mitochondrial phylogeny and provides a ranked classification with an associated quality score. MitoArc uses the phylotree-rcrs@17.3 tree, requests a single highest-ranked assignment, and sets the HaploGrep heteroplasmy-level parameter to 0.9.

Haplogroup classification is attempted only when the input VCF describes a full-length 16,569-bp human mitochondrial reference. Inputs that do not meet this requirement are reported as unavailable rather than classified against an incompatible reference representation. For each caller, MitoArc records the highest-ranked haplogroup, HaploGrep quality value, rank, number of input variants, classification status, software version, and reference tree.

Caller-specific haplogroups are retained separately and are also compared against the same phylogenetic tree. When both assignments are available, they are classified as identical if the assigned haplogroups match, compatible_parent_child if one lies on the ancestral path of the other, or discordant otherwise. No consensus haplogroup is constructed from disagreeing caller assignments.

Sample contamination is assessed independently from each caller-specific canonical VCF using haplocheck 1.3.3. Haplocheck uses mitochondrial phylogenetic information to identify evidence of more than one mitochondrial sequence background within a sample (Weissensteiner et al. 2021). MitoArc retains the native haplocheck output and records the estimated contamination level together with the native contamination status, phylogenetic distance, and sample-coverage measure.

When haplocheck does not provide a numerical contamination estimate, the result is reported as INSUFFICIENT_DATA rather than interpreted as zero contamination. Failed haplocheck analyses are recorded separately as FAILED_QC. Results derived from the GATK and mutserve2 callsets remain separate in the final contamination table and are not averaged or combined into a single estimate.

### 2.10 Functional annotation and consensus generation

Caller-specific variants are annotated using a lightweight MitoArc annotator based on the NCBI RefSeq mitochondrial sequence NC_012920.1, corresponding to the revised Cambridge Reference Sequence (rCRS) (Andrews et al. 1999). The bundled annotation resource contains the 16,569-bp reference sequence and a compact feature table derived from the corresponding NCBI record. Coding sequences, rRNAs, tRNAs, and the control region are included. Protein-coding consequences are interpreted using the vertebrate mitochondrial genetic code (NCBI translation table 2).

Annotation is performed independently for the filtered GATK and normalized mutserve2 callsets. For each alternate allele, MitoArc records the overlapping gene and feature type, region class, descriptive consequence, variant allele fraction (VAF), depth, variant state, caller filter status, and NUMT evidence where available. Simple coding SNVs are translated to determine synonymous, missense, stop-gain, stop-loss, or start-loss consequences. Coding indels are classified as frameshift or in-frame events according to their length change; amino-acid changes are not inferred for these records.

Biological annotation is enabled only when the analyzed mitochondrial sequence is exactly 16,569 bp and is sequence-identical to the bundled NC_012920.1 reference. For incompatible or truncated references, caller metrics are retained, but gene and consequence fields are reported as unavailable.

Variants with VAF ≥0.95 are labeled NEAR_HOMOPLASMIC, whereas variants below this threshold are labeled HETEROPLASMIC; variants without an available VAF are labeled UNKNOWN. The 0.95 threshold follows the convention used for the gnomAD mitochondrial callset (Laricchia et al. 2022). These labels describe allele fraction only and are not clinical classifications.

Caller-specific consensus sequences are generated independently from the canonical GATK and mutserve2 VCFs. Consensus construction begins from the unshifted mitochondrial reference and considers only PASS or unfiltered records. Reference alleles are checked before each edit, overlapping or conflicting edits are rejected, and accepted edits are applied in descending coordinate order to preserve coordinates when indels alter sequence length.

By default, variants with VAF ≥0.95 replace the reference allele, while lower-VAF variants remain represented by the reference base. An optional IUPAC mode can instead represent heteroplasmic SNVs using the appropriate ambiguity code. Heteroplasmic indels remain unchanged because they cannot be represented by a single IUPAC character. Consensus validation records reference mismatches, overlapping edits, conflicting high-VAF alternate alleles, shifted-coordinate records, sequence-length changes, and reference compatibility. GATK and mutserve2 consensus sequences remain separate throughout the analysis.

### 2.11 Reporting and visualization

MitoArc generates machine-readable reporting tables, static figures, an integrated HTML report, and a separate MultiQC report. Reporting is performed after variant calling and interpretation and does not modify caller VCFs, select a preferred caller, or generate a merged callset.

For each sample, a reporting table and matching JSON record collect mitochondrial coverage metrics, the copy-number proxy where available, caller-specific variant counts, haplogroup and contamination results, NUMT evidence, annotation status, consensus validation, analysis thresholds, reference compatibility, and software versions. Separate tables retain per-position mitochondrial depth, caller-specific annotated variants, caller-comparison results, mitochondrial features, and the data used to generate the circular genome view.

Static figures are generated with Matplotlib 3.10.5 (Hunter 2007) and are written as SVG and 180-dpi PNG files. Six figure types are produced: a linear mitochondrial depth profile, coverage-threshold summary, caller-specific VAF landscape, functional-consequence summary, caller-comparison plot, and circular mitochondrial genome view. Coverage values are plotted without smoothing. Variant markers retain caller identity and heteroplasmy state, and variants carrying NUMT-associated evidence are marked separately.

The circular view displays protein-coding genes, rRNAs, tRNAs, and the control region along the mitochondrial genome. Feature strand is shown on separate radial bands, coverage is displayed on a log-scaled radial track, and caller-specific variants occupy separate radii. The linear reference breakpoint is marked explicitly. Circular and biological consequence plots are generated only when the analyzed mitochondrial sequence is compatible with NC_012920.1; otherwise, these panels are reported as unavailable rather than shown as canonical human mtDNA interpretation.

A self-contained HTML report combines the structured results and PNG figures into sections covering sample information, coverage, variants, caller comparison, NUMT evidence, haplogroup, contamination, annotation, consensus status, and reproducibility information. Figures, style information, and table functions are embedded in the report, allowing it to be opened locally without external web resources. The underlying TSV and JSON files remain available separately from the HTML presentation.

Technical quality-control outputs from FastQC, fastp, and samtools are also collected with MultiQC 1.27.1 (Ewels et al. 2016). MultiQC is retained as a separate technical QC report rather than used as the source of MitoArc’s mitochondrial interpretation tables.

### 2.12 Controlled benchmark design

A controlled synthetic benchmark was constructed to examine variant recovery across sequencing depth, allele fraction, sequence position, NUMT-associated mapping context, and caller configuration. The mitochondrial sequence was identical to NC_012920.1 and was represented as a 16,569-bp chrM contig. The benchmark reference also contained 22 synthetic 4-kb autosomal scaffolds and additional mitochondrial-homologous nuclear contigs used for the controlled NUMT experiments. The autosomal scaffolds were included to provide the alignment structure required for full-reference processing; they do not represent a biological whole-genome background.

Eighteen paired-end FASTQ datasets were generated from three independently seeded technical replicates at six target mitochondrial depths: 100×, 250×, 500×, 1000×, 2000×, and 5000×. Each replicate-depth combination used a distinct random seed. Reads were 150 bp with a nominal fragment length of 450 bp. Fragment starts were distributed across the circular mitochondrial sequence and randomized according to the dataset seed. All generated bases were assigned a Phred quality score of 40. No additional sequencing-error model was introduced.

Each dataset contained 36 mitochondrial truth alleles: 24 SNVs and 12 single-base insertions. Six nominal VAF targets were examined: 0.005, 0.015, 0.035, 0.075, 0.25, and 0.75. Each VAF was represented by six fixed archetypes: a non-boundary SNV, a boundary SNV, a non-boundary SNV in a NUMT-like context, a non-boundary SNV in an ambiguous mapping context, a non-boundary insertion, and a boundary insertion. Boundary sites were placed near the ends of the linear mitochondrial reference, and generation of boundary alleles preferentially selected fragments crossing the circular junction.

The number of alternate fragments was determined from the fragments that overlapped each truth position. Consequently, the achieved VAF could differ from the nominal target, particularly at low depth where alternate-fragment counts are discrete. The achieved VAF, rather than the nominal value, was retained as the quantitative truth for VAF-error calculations. Across the 18 datasets, the design contained 648 mitochondrial truth observations.

All datasets were processed through MitoArc using the NUMT comparison mode. Calls were normalized before exact comparison by chromosome, position, reference allele, and alternate allele. Across the seven caller or evidence views retained by the benchmark, this produced 4,536 truth-callset comparisons. The controlled benchmark design is summarized in Table 1.

**Table 1.** Design of the controlled mitochondrial benchmark.

| **Design element** | **Specification** |
|---|---|
| Technical replicates | 3 independently seeded replicates (R1, R2, R3) |
| Target mitochondrial depths | 100×, 250×, 500×, 1000×, 2000×, 5000× |
| Physical benchmark executions | 18 |
| Truth alleles per dataset | 36 |
| Total truth observations | 648 |
| Variant classes per dataset | 24 SNVs and 12 single-base insertions |
| Nominal VAFs | 0.005, 0.015, 0.035, 0.075, 0.25, 0.75 |
| Truth archetypes per VAF | Non-boundary SNV, boundary SNV, NUMT-like SNV, ambiguous-mapping SNV, non-boundary insertion, boundary insertion |
| Read configuration | Paired-end 150 bp |
| Fragment length | 450 bp |
| Mitochondrial reference | NC_012920.1-compatible 16,569-bp sequence |
| Synthetic nuclear reference context | 22 neutral 4-kb scaffolds, 12 challenge homologs, and 12 homologs representing true NUMT-associated context |
| NUMT challenge classes | NUMT_FALSE and AMBIGUOUS |
| Challenge observations | 216 designed NUMT-associated challenges |
| Circular-coordinate evaluation | Boundary and non-boundary truth alleles included |
| Background sequencing-error model | None; reads were generated at constant Q40 |

*Note.* Each nominal VAF was represented by six predefined variant archetypes. Alternate-fragment counts were selected deterministically from the eligible overlapping fragments, so achieved VAF could differ from the nominal value, particularly at lower sequencing depths. The controlled reference was designed for methodological evaluation and does not represent a biological whole-genome sample.

Additional details of benchmark construction and technical replicates are provided in Online Resource
1 (Section S1).

### 2.13 NUMT observability analysis

The NUMT experiment contained both true mitochondrial variants placed in controlled homologous contexts and separate false-signal challenge alleles. For each of the six VAF targets, the truth panel contained one mitochondrial SNV labeled NUMT_LIKE and one labeled AMBIGUOUS_CONTEXT. These alleles remained true mitochondrial variants, but matching mitochondrial-homologous nuclear sequence was added to the synthetic reference to introduce controlled mapping competition. This produced 12 context-labeled true mitochondrial alleles per dataset and 216 such truth observations across the benchmark.

A separate set of 12 challenge alleles was generated for each dataset. At each VAF target, one challenge was classified as NUMT_FALSE and one as AMBIGUOUS. The mitochondrial allele at each challenge position remained the reference allele. Alternate-containing reads were introduced separately and therefore represented false mitochondrial signals if reported as mtDNA variants.

The synthetic nuclear homologs contained a 600-bp mitochondrial-homologous segment with unique synthetic sequence on either side. NUMT_FALSE read pairs combined an alternate-containing mitochondrial-homologous read with a mate derived from unique nuclear sequence. AMBIGUOUS challenge reads were generated from sequence with equivalent mitochondrial and nuclear homology. Challenge-pair counts were determined from the requested VAF and target depth, with a minimum of one pair at the lowest abundance.

Because a designed challenge does not necessarily survive alignment and mitochondrial read extraction, MitoArc recorded a separate observability chain. For each challenge, the benchmark recorded whether its reads were generated, observed after full-reference mapping, associated with a nuclear alternative, and retained during mtDNA extraction. A challenge was considered **observable** only when at least one corresponding fragment entered the mitochondrial analysis path.

NUMT-specific false-positive calculations therefore used the metric scope OBSERVABLE_NUMT_ASSOCIATED_FP. A false signal contributed to this count only if the corresponding designed challenge was observable after mapping and extraction. Designed but unobservable challenges were reported separately rather than counted as caller failures. True mitochondrial retention was assessed independently from the context-labeled truth alleles described above. Additional details of NUMT challenge observability and mapping-context evaluation are provided in Online Resource 1 (Section S2).

### 2.14 Matched GATK circular-coordinate evaluation

A separate matched GATK analysis was performed to isolate the effect of shifted-coordinate processing from differences in calling interval or post-processing. Nine benchmark datasets were selected, comprising all three replicates at 250×, 1000×, and 5000×. Within each dataset, truth alleles at nominal VAFs of 0.015, 0.075, and 0.25 were selected when they had no NUMT challenge. The subset contained boundary and non-boundary SNVs and insertions, giving 108 allele-level truth observations and 216 matched truth-mode observations.

For the standard-only condition, GATK Mutect2 was rerun on the same standard mitochondrial realignment used by MitoArc, but across the complete chrM:1-16569 interval. FilterMutectCalls was then applied using mitochondrial mode. This avoided using the interval-restricted standard branch that forms only one component of MitoArc’s normal standard-plus-shifted procedure.

The resulting standard-only callset was compared with the canonical standard-plus-shifted GATK callset from the same dataset. Both were normalized to the same mitochondrial reference and compared by exact allele identity. Recall and absolute VAF error were evaluated separately for boundary and non-boundary SNVs and insertions. Additional details of the matched circular-coordinate evaluation are provided in Online Resource 1 (Section S4).

### 2.15 mutserve2 indel-configuration analysis

The primary controlled panel contained 216 single-base insertion truth observations across all depths and replicates. To determine whether missing insertion calls arose during normalization or earlier in the calling process, each truth observation was traced through the native mutserve2 VCF, the normalized canonical VCF, and the benchmark comparison layer. This analysis was performed without changing the frozen MitoArc v1.0.0 mutserve2 command, in which the beta insertion and deletion options were disabled.

An additional positive-control dataset was generated to test the frozen configuration with direct read support for both an insertion and a deletion. The control contained a single-base insertion at mitochondrial position 8000 and a single-base deletion at position 9000, each at a nominal VAF of 0.25 and a target depth of 1000×. An unrelated SNV was included to ensure that the caller produced a non-empty VCF. Alternate- and reference-supporting fragments were counted directly from the aligned BAM using their CIGAR representation.

The positive-control alignment was analyzed with both GATK Mutect2 and mutserve2. mutserve2 was run with the same v1.0.0 command used in the primary benchmark, without enabling its insertion or deletion options. The positive control was used only to identify the stage at which indel observations were lost and was not included in the primary caller-performance totals.

### 2.16 HG002 real-data validation

End-to-end validation was also performed with the Genome in a Bottle (GIAB) HG002/NA24385 reference sample (Zook et al. 2019). A publicly available Illumina HiSeq 300× BAM aligned to GRCh38 (HG002.GRCh38.300x.bam; MD5: a2a85243fba525d485a6ba89d73b46a9) was accessed through the Genome in a Bottle alignment index. Because the complete source BAM was approximately 601 GB, MitoArc validation used indexed regional retrieval rather than downloading or processing the complete alignment.

The reduced BAM retained all alignments overlapping chrM together with three 100-kb nuclear intervals: chr1:600001-700000, chr2:10000001-10100000, and chr17:22500001-22600000. The chr1 interval was included to retain a mitochondrial-homologous nuclear context, while the chr2 and chr17 intervals supplied additional genuine nuclear alignment context. All chrM alignments were retained without subsampling. Regional extraction did not alter read sequence, base quality, mapping quality, CIGAR strings, or alignment tags. The resulting BAM was coordinate sorted and indexed with samtools 1.20.

The matching GIAB GRCh38 no-alt analysis reference was used for processing. Its mitochondrial sequence was confirmed to be identical to NC_012920.1. Because only selected nuclear intervals were retained, the input was treated as PARTIAL_NUCLEAR_CONTEXT, not as complete whole-genome data.

The reduced HG002 BAM was processed through the pre-aligned MitoArc path, including mitochondrial extraction, coverage analysis, circular-coordinate processing, both variant callers, NUMT evidence assessment, haplogroup and contamination analysis, annotation, consensus generation, and reporting. No mtDNA truth set was assigned to this experiment. HG002 was therefore used to assess end-to-end operation on real human sequencing data rather than to estimate variant-calling accuracy.

### 2.17 Computational resource measurement

Resource use was recorded for each of the 18 controlled benchmark executions. Benchmark scheduling was restricted to one active workflow process at a time using a Nextflow queue size of one and maxForks = 1. The designated primary successful execution for each dataset was used for resource scaling; later resume-validation runs were not included.

Wall-clock time was measured by the benchmark harness from Nextflow invocation to workflow completion. The size of the Nextflow work directory was sampled at 30-second intervals and again at workflow completion to estimate peak temporary storage. Final output size was calculated from the corresponding results directory.

CPU use and memory were obtained from the Nextflow execution trace. CPU time was estimated for submitted processes from process real time and reported CPU utilization and was converted to CPU-hours. Peak resident-set size was defined as the maximum reported peak RSS among submitted processes. Cached processes were excluded from CPU and memory calculations.

These measurements apply to the complete physical MitoArc execution, which generated both caller outputs and the NUMT comparison branches. They should therefore not be interpreted as separate GATK or mutserve2 runtime measurements. Resource values are also dependent on the execution environment and are reported as observed benchmark measurements rather than general hardware requirements. Additional resource measurements and the software-version manifest are provided in Online Resource 1 (Section S5).

### 2.18 Statistical and benchmark analysis

Variant calls were normalized before comparison with the truth panel. A true positive (TP) required an exact normalized match of chromosome, position, reference allele, and alternate allele. A truth allele without an active matching call was classified as a false negative (FN). An active call not present in the mitochondrial truth panel was classified as a false positive (FP). Caller disagreement alone was not considered an error.

Two false-positive scopes were retained because they answer different questions. TOTAL_EXACT_TRUTH_FP includes all active calls not matching the controlled mitochondrial truth and was used for the pooled GATK and mutserve2 comparison. OBSERVABLE_NUMT_ASSOCIATED_FP is restricted to designed NUMT or ambiguous challenge alleles that remained observable after full-reference mapping and mitochondrial extraction and was used for the NUMT-strategy analysis.

Precision, recall, and F1 score were calculated as:

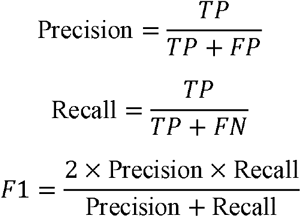

For recovered truth alleles, absolute VAF error was calculated as the absolute difference between the caller-reported VAF and the achieved synthetic VAF. Relative VAF error was calculated by dividing the absolute error by the achieved VAF. VAF strata were assigned from the achieved rather than nominal allele fraction: <0.01, 0.01–<0.02, 0.02–<0.05, 0.05–<0.10, 0.10–0.50, and >0.50.

Benchmark outcomes were summarized by depth, VAF, variant class, boundary class, NUMT context, caller, and NUMT strategy. Replicate-level summaries included means, medians, standard deviations, ranges, and interquartile values where applicable. For depth-specific VAF error, the mean absolute error was calculated within each replicate and then summarized across the three replicates.

The three benchmark replicates were stochastic technical replicates generated at fixed truth positions; they were not independent biological samples. The analysis was therefore treated as descriptive. No inferential hypothesis tests or *p*-values were applied, and the benchmark was not used to define an optimal sequencing depth or a universal ranking between callers.

## 3. Results

### 3.1 MitoArc supports FASTQ and pre-aligned inputs through a shared mtDNA analysis path

MitoArc was organized around three input routes for paired-end FASTQ, coordinate-sorted BAM, and CRAM data (Fig. 1a). FASTQ input passes through read quality control, preprocessing, and full-reference alignment, whereas BAM and CRAM inputs enter through pre-aligned input validation. The three routes then converge on the same downstream mtDNA analysis, including mitochondrial read extraction, coverage assessment, standard and shifted-coordinate processing, variant calling, NUMT evidence assessment, haplogroup and contamination analysis, functional annotation, consensus generation, and reporting. GATK and mutserve2 outputs remain separate throughout the workflow.

**Fig. 1.**
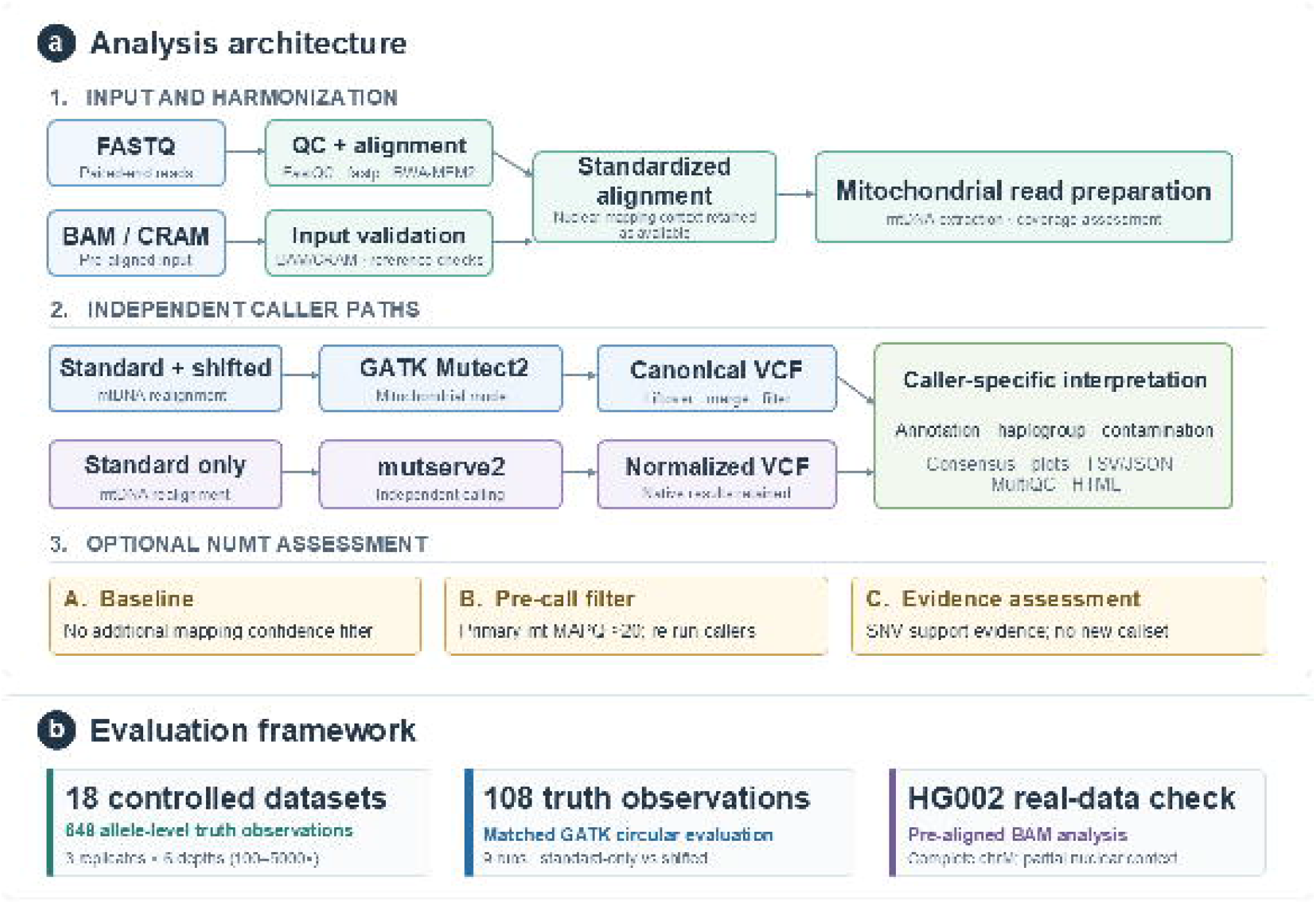
MitoArc analysis architecture and evaluation framework. (a) Paired-end FASTQ data undergo quality control, preprocessing, and full-reference alignment, while validated BAM and CRAM inputs enter through the pre-aligned route. Both routes converge on a standardized alignment before mitochondrial read extraction and coverage assessment. GATK Mutect2 uses standard and shifted mitochondrial reference alignments, with shifted calls returned to canonical coordinates before combination and filtering. mutserve2 operates independently on the standard mitochondrial alignment. Optional NUMT analysis comprises baseline calling (Strategy A), pre-calling mapping-confidence filtering followed by repeat variant calling (Strategy B), and supporting-fragment evidence classification without generating an additional callset (Strategy C). Caller-specific results remain separate during interpretation and reporting. (b) Evaluation includes 18 controlled datasets comprising three technical replicates at six sequencing depths (100×–5000×), with 648 allele-level truth observations; a matched GATK circular-coordinate analysis involving 108 truth observations from nine datasets; and reduced HG002 real-data analysis with partial nuclear context to assess end-to-end workflow operation

The controlled benchmark exercised the FASTQ route across 18 independently generated datasets comprising three replicates at six sequencing depths (Fig. 1b). All 18 benchmark executions completed successfully and produced the expected caller, NUMT, interpretation, and reporting outputs. The real-data validation independently exercised the pre-aligned BAM route using HG002 and completed the full downstream workflow, including both variant callers, NUMT evidence analysis, haplogroup assignment, contamination assessment, functional annotation, consensus generation, MultiQC, static figures, and the self-contained HTML report. The CRAM route was tested with dedicated input and reference-validation fixtures but was not used for the publication-scale benchmark or HG002 experiment.

Process reuse was also checked after successful execution. An identical resumed HG002 analysis reused all 61 completed processes without submitting new tasks. Benchmark resume checks at 100×, 1000×, and 5000× similarly reused all 67 completed processes for each tested run without new task submission. These checks confirmed stable Nextflow caching for the evaluated executions.

### 3.2 Mapping-context filtering removes observable synthetic NUMT-associated false positives at a cost to variant recovery

The effect of mapping-context filtering was evaluated by comparing GATK Strategy A, representing the baseline calling path, with Strategy B, in which read names were retained only when supported by a primary mitochondrial alignment meeting the mapping-quality threshold. The analysis was performed across all 18 controlled benchmark datasets (Fig. 2).

**Fig. 2.**
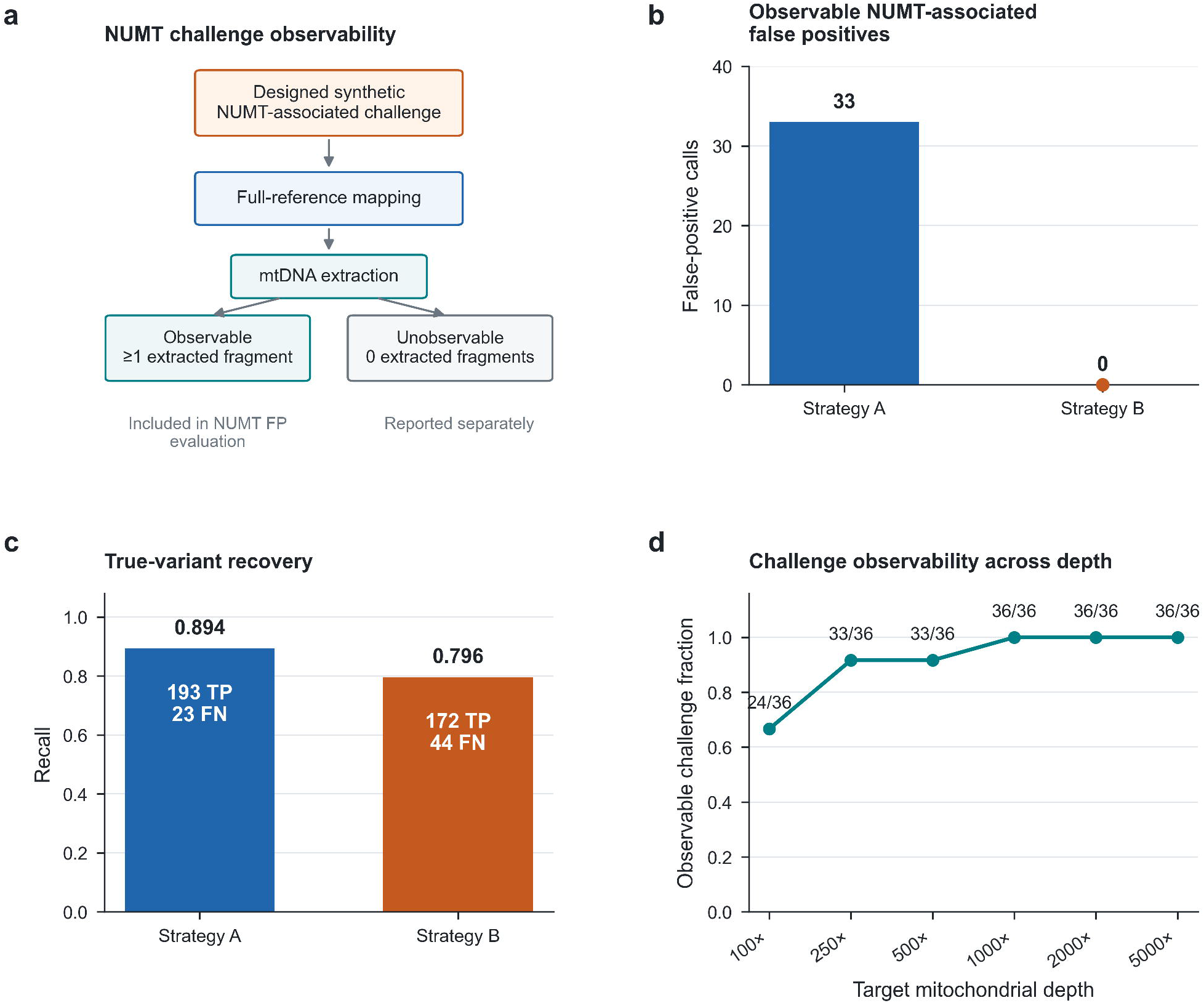
Mapping-context filtering reduces observable synthetic NUMT-associated false-positive calls at a cost to true-variant recovery. (a) Synthetic NUMT-associated challenges were tracked through full-reference mapping and mtDNA extraction; only challenges with at least one extracted fragment were considered observable for NUMT-associated false-positive evaluation. (b) Strategy B eliminated the 33 observable NUMT-associated false-positive calls detected under Strategy A. (c) This reduction was accompanied by lower true-variant recovery, with recall decreasing from 0.894 to 0.796. (d) Challenge observability increased with sequencing depth, reaching 36 of 36 challenges from 1000× onward

The NUMT experiment contained 216 designed challenge instances across the complete benchmark. Not every challenge remained observable after full-reference mapping and mitochondrial read extraction. Across the six depth levels, 198 of 216 challenge instances entered the mitochondrial analysis path. Observability increased with depth: 24 of 36 challenge instances were observable at 100×, 33 of 36 at both 250× and 500×, and all 36 at 1000×, 2000×, and 5000× (Online Resource 1, Fig. S3). Designed challenges that did not enter the mitochondrial analysis path were excluded from the NUMT-associated false-positive count rather than treated as successfully removed.

Using this observability-aware comparison, Strategy A recovered 193 true mitochondrial observations, produced 33 observable NUMT-associated false-positive calls, and missed 23 true mitochondrial observations. This corresponded to a precision of 0.854, recall of 0.894, and F1 score of 0.873. Strategy B produced no observable NUMT-associated false positives, while recovering 172 true mitochondrial observations and missing 44. Precision increased to 1.000, recall decreased to 0.796, and F1 score was 0.887. Mapping-context filtering therefore removed all 33 observable NUMT-associated false-positive calls in this benchmark, but 21 fewer true mitochondrial observations were recovered. The pooled Strategy A and Strategy B performance metrics are summarized in Table 2.

**Table 2.** Effect of mapping-context filtering on observable synthetic NUMT-associated false positives.

| **Strategy** | **TP** | **Observable NUMT-associated FP** | **FN** | **Precision** | **Recall** | **F1** |
|---|---|---|---|---|---|---|
| Strategy A | 193 | 33 | 23 | 0.854 | 0.894 | 0.873 |
| Strategy B | 172 | 0 | 44 | 1.000 | 0.796 | 0.887 |

*Note.* False positives in this table use the OBSERVABLE\_NUMT\_ASSOCIATED\_FP definition. Only designed NUMT-associated challenges with at least one fragment retained after mapping and mtDNA extraction were eligible false-positive opportunities. Of 216 designed challenges, 198 were observable. Because TP/FN and observable NUMT-associated FP derive from separate designed opportunity sets, precision and F1 are specific to the composition of this synthetic benchmark and are interpreted descriptively.

The same pattern was present in each replicate. Strategy A produced 11 observable NUMT-associated false positives in each of the three replicates, whereas Strategy B produced none. True-positive counts changed from 65 to 58 in replicate 1 and from 64 to 57 in both replicates 2 and 3. The reduction in false-positive calls was therefore accompanied by lower true-variant recovery in each independently generated replicate.

Strategy C was not included as a primary mixed-variant performance comparison. Its fragment-level support analysis is restricted to SNVs, causing non-SNV alleles to enter an unresolved-support category. Targeted diagnostic tracing confirmed that supporting reads for representative unresolved indels were present in the mitochondrial alignments; their unresolved status resulted from the defined scope of the Strategy C classifier rather than absence of supporting reads (Online Resource 1, Fig. S2).

### 3.3 Shifted-coordinate processing shows no incremental boundary-specific recovery in the matched GATK benchmark

The effect of shifted-coordinate processing was examined in a matched GATK analysis using nine benchmark datasets spanning three replicates and mitochondrial depths of 250×, 1000×, and 5000×. The analysis included 108 allele-level truth observations at nominal VAFs of 0.015, 0.075, and 0.25, with boundary and non-boundary SNVs and insertions represented in each comparison. Standard-only and standard-plus-shifted callsets were subjected to equivalent calling, filtering, and normalization before comparison (Online Resource 1, Fig. S1).

All 108 allele-level calling results were identical between the two processing modes. Boundary SNV recall was 0.778 in both modes, with 21 true positives and six false negatives. Boundary insertion recall was 1.000 in both modes, with all 27 truth observations recovered. Non-boundary SNV recall was 1.000 in both modes, while non-boundary insertion recall was 0.926, with 25 of 27 truth observations recovered. No recall difference was observed between standard-only and standard-plus-shifted processing for any of the four variant classes.

Caller-reported VAFs were also identical between processing modes for every recovered truth observation. The matched experiment therefore found no incremental effect of shifted-coordinate processing on either variant recovery or VAF estimation within the evaluated boundary fixture.

### 3.4 Sequencing depth influences variant recovery, VAF accuracy, and computational demand

Variant recovery was evaluated across mitochondrial depths of 100×, 250×, 500×, 1000×, 2000×, and 5000× using three independently generated replicates at each depth (Fig. 3). For GATK Strategy A, mean recall increased from 0.741 at 100× to 0.806 at 250×, 0.843 at 500×, 0.880 at 1000×, 0.889 at 2000×, and 0.917 at 5000×. The increase was largest over the lower depth range, while the change above 1000× was smaller.

**Fig. 3.**
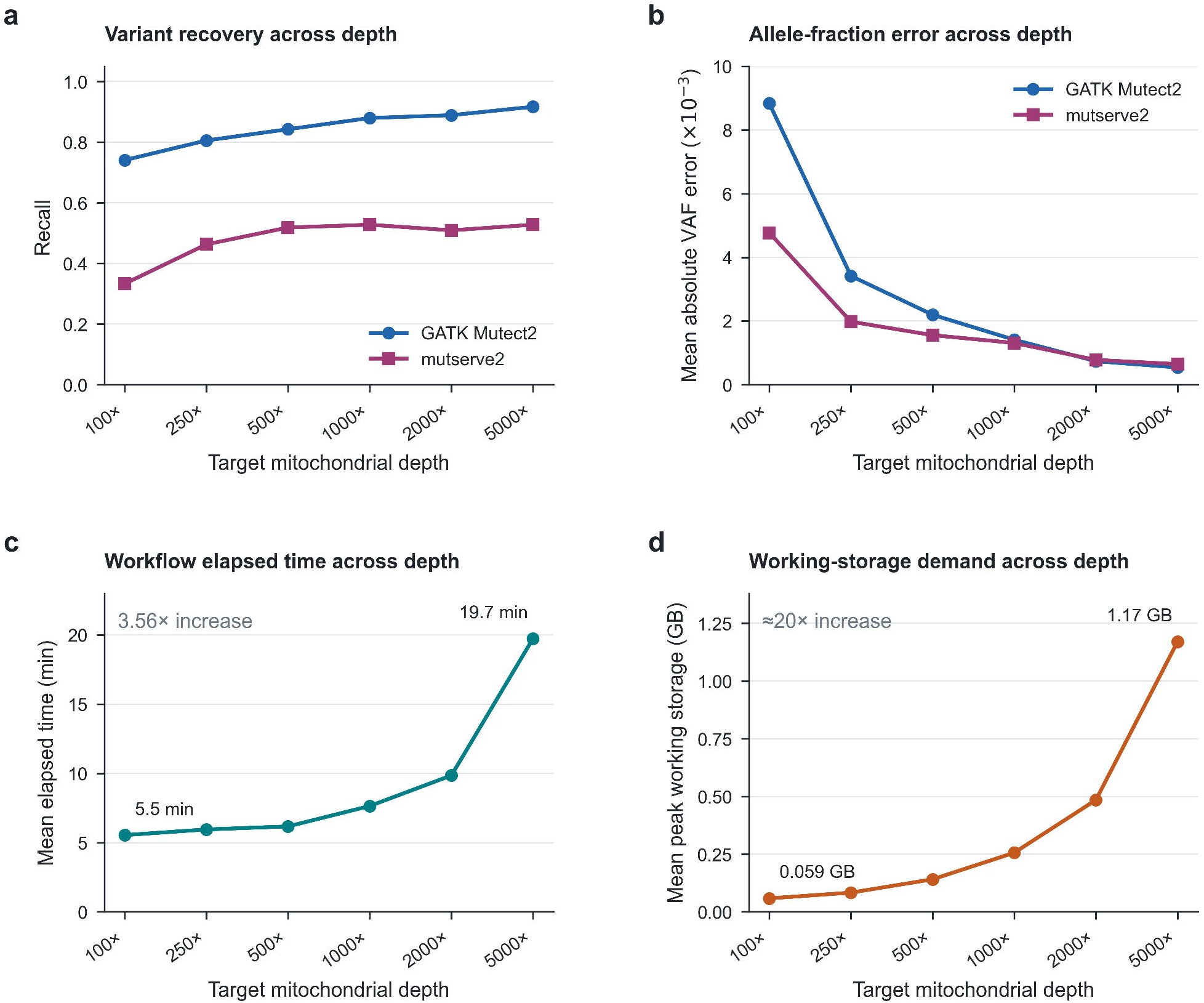
Sequencing depth influences variant recovery, allele-fraction error, and computational demand. (a) Recall across sequencing depth for GATK Mutect2 and mutserve2 Strategy A callsets. (b) Mean absolute variant allele fraction (VAF) error among recovered truth observations decreased with increasing depth for both callers; values should be interpreted alongside caller-specific recovery because the recovered truth subsets differed. (c) Mean workflow elapsed time increased from approximately 5.5 min at 100× to 19.7 min at 5000×. (d) Mean peak working-directory storage increased from approximately 0.059 GB to 1.17 GB across the same depth range. Resource measurements represent complete workflow executions and are not caller-specific

mutserve2 showed a different depth response under the evaluated configuration. Mean recall increased from 0.333 at 100× to 0.463 at 250× and 0.519 at 500×. Recall was 0.528 at 1000×, 0.509 at 2000×, and 0.528 at 5000×. Variant recovery therefore changed little beyond 500× in this benchmark.

Allele-fraction estimation improved with increasing depth for both callers. Mean absolute VAF error for GATK decreased from 0.00884 at 100× to 0.00342 at 250×, 0.00220 at 500×, 0.00141 at 1000×, 0.00074 at 2000×, and 0.00054 at 5000×. For mutserve2, the corresponding values were 0.00477, 0.00198, 0.00156, 0.00131, 0.00078, and 0.00065. Low-frequency achieved VAFs were more strongly quantized at lower sequencing depths, particularly at 100×, because individual alternate-supporting fragments represented a larger fraction of the available coverage.

Increasing mitochondrial depth also increased computational demand. Mean workflow runtime rose from 333 s at 100× to 1184 s at 5000×, a 3.56-fold increase. Mean peak working-directory storage increased from 58.5 MB to 1.17 GB over the same range, corresponding to a 20-fold increase. Mean CPU use increased from 0.102 CPU-hours per run at 100× to 0.237 CPU-hours at 5000×. Peak resident memory did not increase at the same rate and reached a maximum observed value of 1.4 GB.

The benchmark therefore showed continued improvement in GATK recall and in VAF estimation with increasing depth, while mutserve2 recall changed little after 500× under the tested configuration. At the same time, temporary storage increased more rapidly than elapsed runtime. The evaluated depth range does not identify a single optimal sequencing depth because the balance between recovery, VAF accuracy, and computational cost depends on the analytical objective.

### 3.5 Variant-caller behavior differs under a shared truth framework and depends on configuration

GATK Mutect2 and mutserve2 were compared against the same 648 truth observations across the 18 controlled benchmark datasets using their Strategy A callsets (Fig. 4). GATK recovered 548 truth observations, produced 33 false-positive calls, and missed 100 truth observations. This corresponded to a precision of 0.943, recall of 0.846, and F1 score of 0.892. mutserve2 recovered 311 truth observations, produced 17 false-positive calls, and missed 337 truth observations, corresponding to a precision of 0.948, recall of 0.480, and F1 score of 0.637. The pooled caller-performance metrics are summarized in Table 3.

**Fig. 4.**
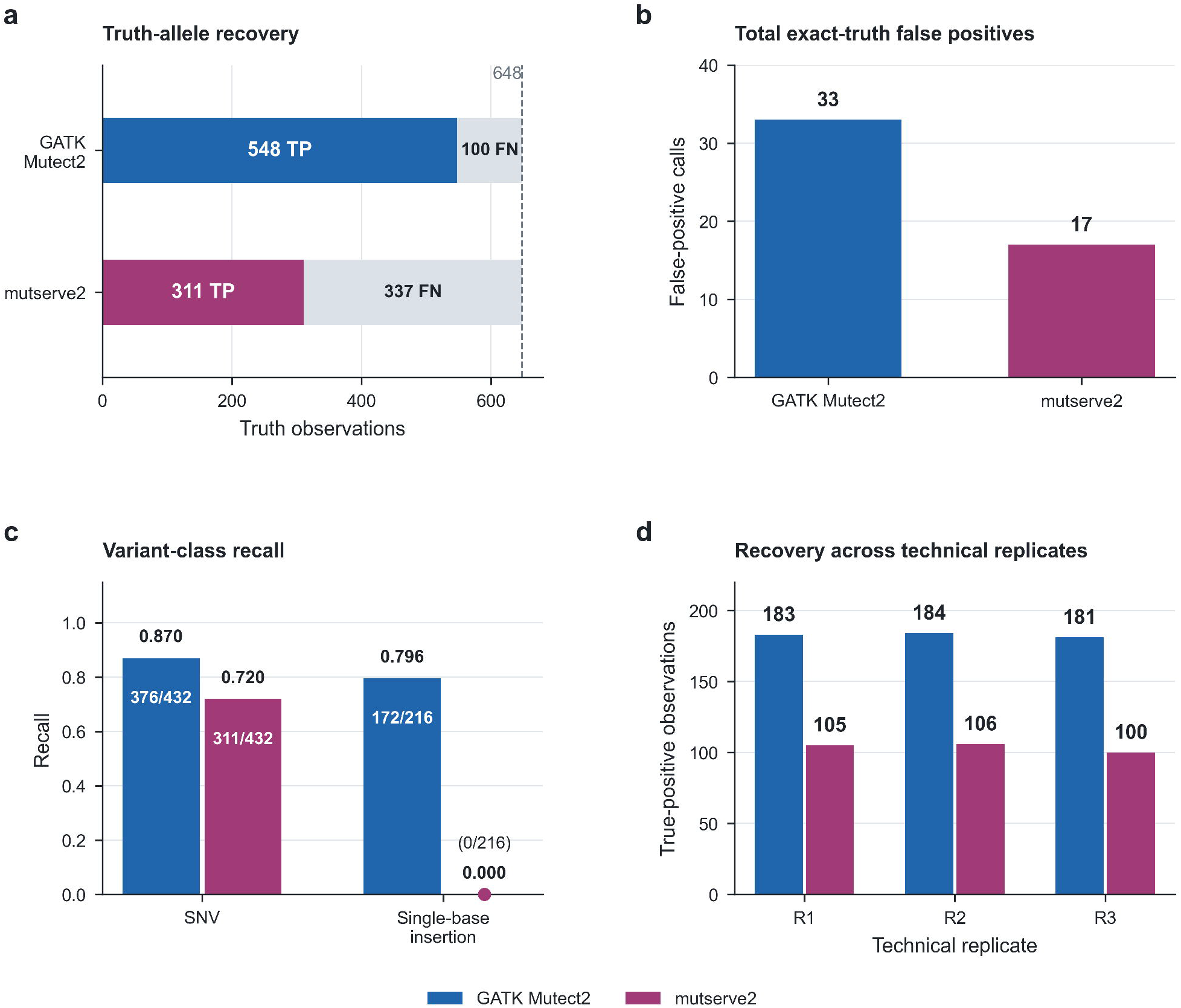
GATK Mutect2 and mutserve2 show distinct recovery patterns under a shared truth framework. (a) Recovery of the 648 allele-level truth observations by each caller. (b) Total exact-truth false-positive calls using the TOTAL_EXACT_TRUTH_FP definition. (c) Recall differed by variant class: both callers recovered SNVs, whereas mutserve2 emitted no truth insertions under the frozen v1.0.0 configuration. This insertion result is configuration-specific because the evaluated mutserve2 invocation did not enable its beta insertion and deletion modes. (d) Caller-specific true-positive recovery across the three independently generated technical replicates

**Table 3.** Controlled benchmark performance of GATK Mutect2 and mutserve2 under a shared truth framework.

| **Caller** | **TP** | **FP** | **FN** | **Precision** | **Recall** | **F1** | **SNV recall** | **Insertion recall** |
|---|---|---|---|---|---|---|---|---|
| GATK Mutect2 | 548 | 33 | 100 | 0.943 | 0.846 | 0.892 | 0.870 (376/432) | 0.796 (172/216) |
| mutserve2 | 311 | 17 | 337 | 0.948 | 0.480 | 0.637 | 0.720 (311/432) | 0.000 (0/216) |

*Note*. Performance was evaluated against 648 exact normalized truth observations using each caller's Strategy A callset. FP denotes TOTAL\_EXACT\_TRUTH\_FP and is therefore distinct from the NUMT-associated false-positive definition used in Table 2. The mutserve2 insertion result is specific to the frozen MitoArc v1.0.0 configuration, in which mutserve2 beta insertion and deletion modes were not enabled.

The callers also differed by variant class. Of the 432 SNV truth observations, GATK recovered 376, giving an SNV recall of 0.870. mutserve2 recovered 311, corresponding to an SNV recall of 0.720. GATK additionally recovered 172 of the 216 single-base insertion truth observations, giving an insertion recall of 0.796. No truth insertions were emitted by mutserve2 under the evaluated configuration.

The absence of mutserve2 insertion calls was examined separately to distinguish caller configuration from downstream processing. All 216 insertion truth observations were traced from native mutserve2 output through VCF conversion and benchmark normalization. No matching insertion was present in the native caller output, and none was subsequently lost during normalization. In a separate positive-control experiment at approximately 1000× depth, the frozen mutserve2 configuration did not emit a known insertion or deletion despite direct read support, while an unrelated SNV control was called successfully. The evaluated mutserve2 invocation had its beta insertion and deletion options disabled. The zero-insertion result therefore describes the configuration tested in MitoArc v1.0.0 and does not indicate that mutserve2 is generally unable to call indels.

Among recovered truth observations, mean absolute VAF error was 0.00267 for GATK and 0.00166 for mutserve2. This comparison applies only to variants recovered by each caller; the two error means were calculated from different sets of truth observations because the callers differed substantially in recall.

Caller recovery was similar across the three independently generated replicates. GATK recovered 183, 184, and 181 truth observations in replicates 1, 2, and 3, respectively. The corresponding mutserve2 totals were 105, 106, and 100. Differences between callers were therefore present across all three replicates rather than being attributable to a single generated dataset.

The false-positive totals in this comparison use the complete exact-truth definition and should not be confused with the NUMT-specific metric used in Section 3.2. The 17 mutserve2 false-positive calls were truth-external calls outside the observable NUMT-associated false-positive category; mutserve2 had no observable NUMT-associated false positives in the Strategy A NUMT analysis. Caller performance therefore depended both on the metric being considered and on the caller configuration used in the benchmark.

### 3.6 HG002 real data support end-to-end workflow operation

MitoArc was evaluated on a reduced alignment derived from the Genome in a Bottle HG002/NA24385 Illumina GRCh38 dataset (Fig. 5). The input retained all chrM-overlapping alignments together with selected nuclear regions on chromosomes 1, 2, and 17. Because the nuclear component was intentionally incomplete, the alignment was classified as PARTIAL_NUCLEAR_CONTEXT and was not treated as a whole-genome accuracy benchmark.

**Fig. 5.**
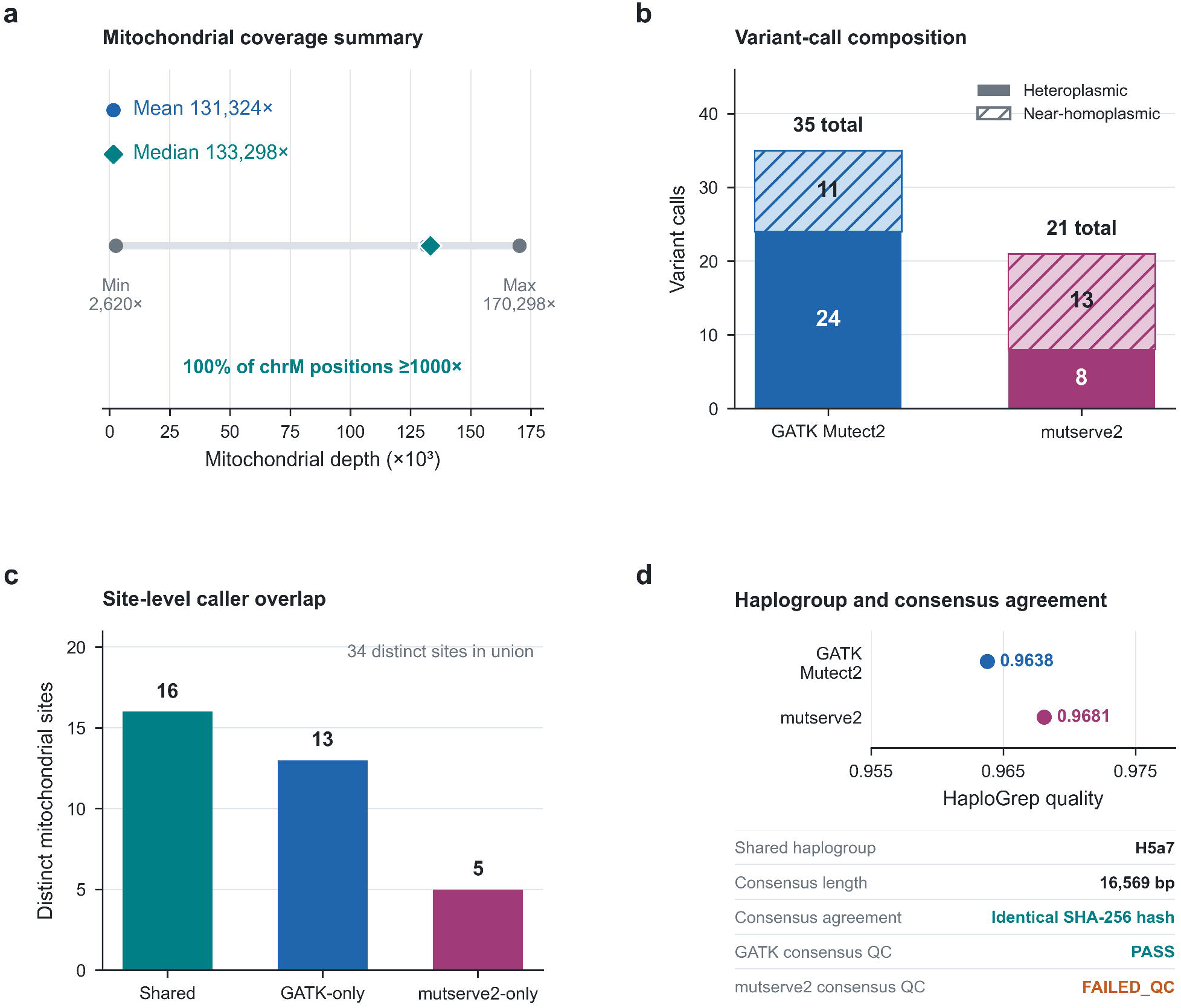
HG002 real data support end-to-end MitoArc workflow operation. (a) Mitochondrial coverage in the reduced HG002 alignment, with mean and median depths of 131,324× and 133,298× and complete coverage above 1000×. (b) Caller-specific heteroplasmic and near-homoplasmic variant-call composition. (c) Site-level overlap comprised 16 shared, 13 GATK-only, and five mutserve2-only sites. (d) Both callsets produced haplogroup H5a7 with similar HaploGrep quality scores, and their 16,569-bp consensus sequences had identical SHA-256 hashes despite different consensus QC statuses. Because the HG002 input had reduced nuclear context and no assigned mitochondrial truth set, this analysis represents integration validation rather than a truth-based accuracy benchmark

Mitochondrial coverage was high across the complete 16,569-bp sequence. Mean depth was 131,324.42× and median depth was 133,298×, with a minimum of 2,620× and maximum of 170,298×. Every mitochondrial position exceeded 1×, 10×, 100×, 500×, and 1,000× coverage. Autosomal depth and the sequencing-derived mtDNA copy-number proxy were reported as unavailable because the reduced alignment did not contain sufficient mapped autosomal context.

GATK reported 35 mitochondrial variant calls, including 24 heteroplasmic and 11 near-homoplasmic calls, with VAFs ranging from approximately 0.0064 to 1.0. mutserve2 reported 21 mitochondrial variant calls, including eight heteroplasmic and 13 near-homoplasmic calls, with VAFs ranging from 0.012 to 1.0. At the site level, 16 sites were shared between the callers, while 13 were GATK-only and five were mutserve2-only.

Both caller-specific callsets produced the same rank-1 haplogroup assignment, H5a7. HaploGrep quality scores were 0.9638 for the GATK callset and 0.9681 for the mutserve2 callset. Haplocheck did not return a supported numerical contamination estimate for either caller, and both results were therefore retained as INSUFFICIENT_DATA.

Functional annotation was available for both callsets across protein-coding genes, rRNAs, tRNAs, and the control region. The GATK consensus passed validation and produced a 16,569-bp mitochondrial sequence containing 11 near-homoplasmic substitutions and no change in sequence length. The mutserve2-derived consensus had the same length and SHA-256 sequence hash as the GATK consensus and therefore contained the same 11 substitutions. Its validation status nevertheless remained FAILED_QC because one multi-allelic conflict was detected during consensus construction.

The HG002 integration results are summarized in Table 4.

**Table 4.**
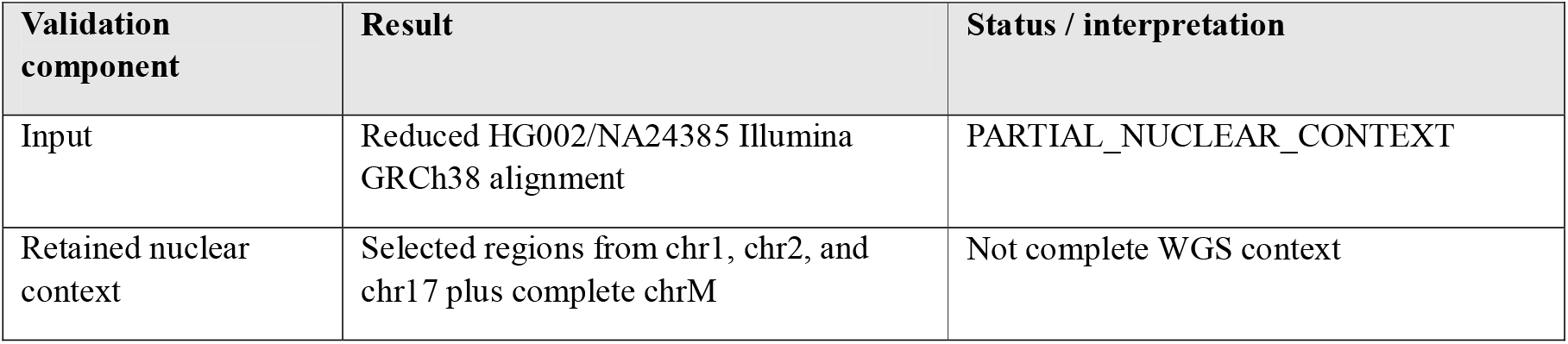
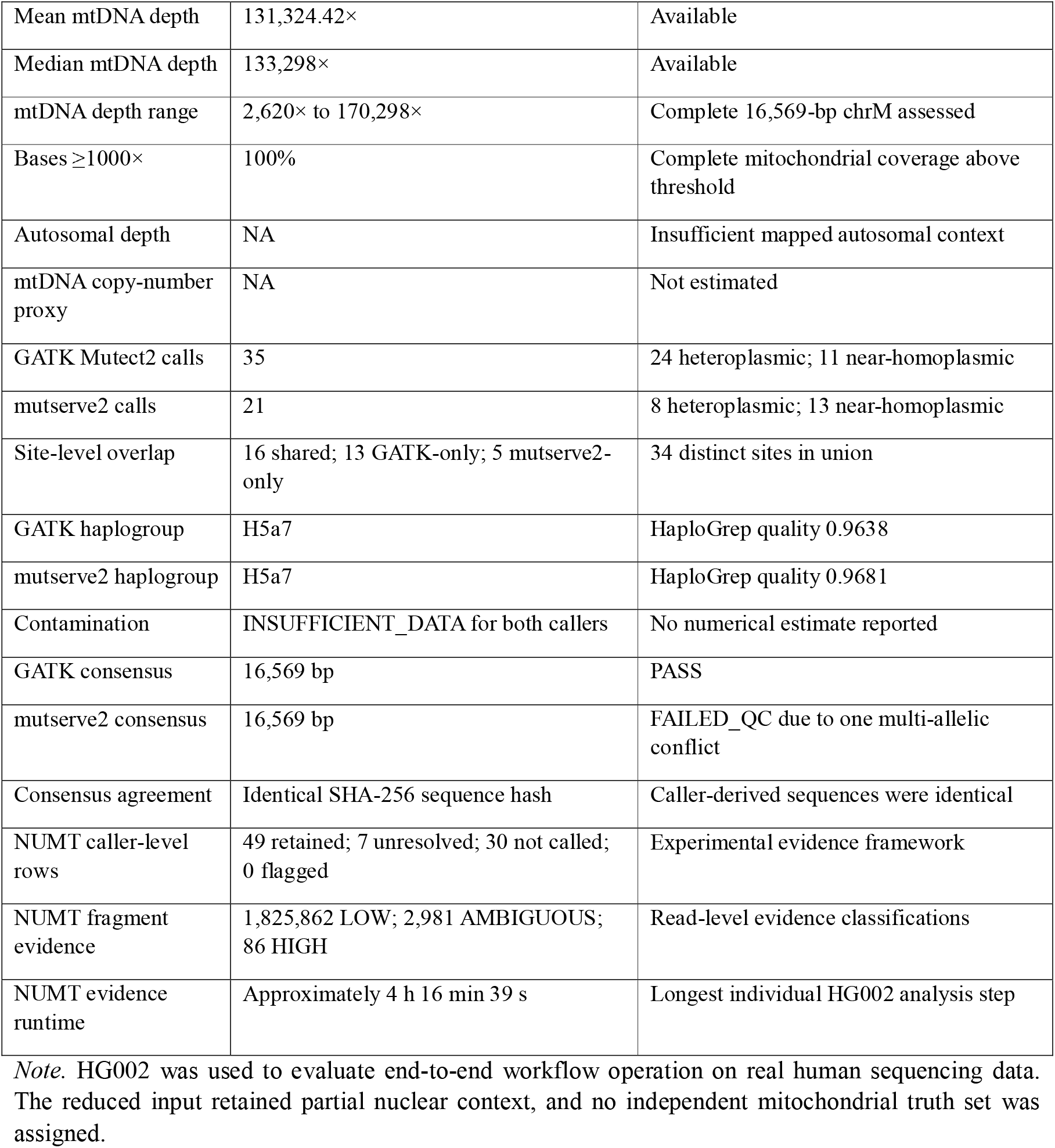
Summary of HG002 real-data integration validation.

NUMT evidence generation was the longest individual step in the HG002 analysis, requiring approximately 4 h 16 min 39 s. The final read-level evidence contained 1,825,862 LOW, 2,981 AMBIGUOUS, and 86 HIGH classifications. At the caller-row level, 49 variants were retained, seven had unresolved support, 30 were not called in the compared context, and none met the criteria for FLAGGED_NUMT_EVIDENCE.

The HG002 analysis therefore exercised the pre-aligned input path through mtDNA extraction, circular-coordinate processing, both callers, NUMT evidence assessment, haplogroup and contamination analysis, functional annotation, consensus construction, visualization, and reporting on real human sequencing data. Because the input contained reduced nuclear context and no independent mtDNA truth set, this experiment was not used to estimate sensitivity, specificity, complete-WGS NUMT performance, contamination accuracy, mtDNA copy-number accuracy, or relative caller accuracy.

## 4. Discussion

MitoArc was developed around the premise that human short-read mtDNA analysis depends not only on the choice of software, but also on decisions about mapping context, reference representation, sequencing depth, and caller configuration. The controlled benchmark supports this view. Requiring confident primary mitochondrial mapping removed the observable synthetic NUMT-associated false-positive signals in the tested data, but also reduced recovery of true mitochondrial observations. Increasing sequencing depth improved GATK variant recovery and allele-fraction estimation, although the gains became smaller at higher depths while computational requirements continued to increase. GATK Mutect2 and mutserve2 showed different recovery patterns under the same truth panel, and the matched GATK circular-coordinate experiment found no additional boundary-specific recovery from shifted processing under the conditions tested. Together, these results argue against treating any single analytical choice as uniformly beneficial across mtDNA analyses.

### 4.1 NUMT filtering involves a trade-off between false-positive suppression and true-variant recovery

NUMTs remain one of the principal sources of ambiguity in short-read mitochondrial analysis. Their sequence similarity to mtDNA can produce both directions of mapping error: nuclear-derived reads can contribute apparent mitochondrial variants, while genuine mitochondrial reads can preferentially map to homologous nuclear regions. Maude et al. (2019) demonstrated this second effect systematically, showing that mitochondrial alleles can lose mitochondrial alignment coverage when the alternate sequence increases similarity to a NUMT, with consequent underestimation or loss of heteroplasmic variants. Large-scale gnomAD analysis likewise showed that NUMT-derived false-positive calls are concentrated particularly among low-VAF observations and are influenced by mitochondrial copy number (Laricchia et al. 2022).

The MitoArc benchmark demonstrates the consequence of this ambiguity directly. Strategy B eliminated all 33 observable synthetic NUMT-associated false-positive calls detected under Strategy A, but recovered 21 fewer true mitochondrial observations and reduced recall from 0.894 to 0.796. The result therefore does not support treating stricter mitochondrial mapping-confidence filtering as an unqualified improvement. A filter that increases confidence in mitochondrial mapping can also exclude genuine mtDNA signal when mitochondrial and nuclear sequence contexts are difficult to distinguish.

Existing mtDNA workflows address this problem in different ways. MToolBox incorporates nuclear realignment during NumtS filtering, whereas MitoHPC attempts to recover mitochondrial reads from selected nuclear regions before subsequent mitochondrial analysis (Battle et al. 2022; Calabrese et al. 2014). The gnomAD mitochondrial pipeline combines whole-genome mapping information, mitochondrial copy-number criteria, and variant-level filters to reduce NUMT-associated artifacts (Laricchia et al. 2022). These approaches are not directly equivalent to MitoArc Strategy B, and the present benchmark was not designed as a head-to-head comparison between NUMT methods. Instead, the controlled result shows why the effect of any filtering rule should be evaluated against both false-positive reduction and loss of true variants.

Challenge observability was also dependent on sequencing depth. Some engineered nuclear-associated signals were generated and mapped but never entered the mitochondrial extraction path, particularly at lower depth. Counting these cases as successfully removed would have overstated the effect of the downstream NUMT strategy. Separating generation, mapping, extraction, and calling therefore provided an important denominator for the NUMT experiment. This distinction is especially relevant for synthetic benchmarking, where absence from a final callset can result from several earlier stages rather than from the filter being evaluated.

The limits of reference-based NUMT handling should also be recognized. Recent long-read pangenome analysis identified both fixed and polymorphic NUMTs, including events not represented by a single conventional human reference, and reported improved detection using a pangenome-based approach (Fu et al. 2026). MitoArc does not resolve this wider problem. Its NUMT strategies evaluate mapping evidence against the supplied reference and should therefore be interpreted as short-read, reference-dependent evidence rather than complete determination of mitochondrial or nuclear origin.

### 4.2 The benefit of shifted-coordinate processing depends on the conditions being tested

Circular-reference handling is well motivated biologically. Linear representation of the mitochondrial genome creates an artificial breakpoint, and shifted or circularized references are used in established mtDNA methods to recover reads and variants affected by this boundary. The gnomAD mitochondrial workflow used an 8,000-bp shift and called the control-region boundary from the shifted alignment, while MitoHPC reported improved coverage around the ends of chrM after mitochondrial realignment to a circularized reference (Battle et al. 2022; Laricchia et al. 2022).

A direct comparison between the original standard-only source and the canonical standard-plus-shifted callset would have been confounded by differences in downstream processing. We therefore performed a matched GATK experiment in which calling, filtering, and normalization were held constant between processing modes. Across 108 allele-level truth observations, standard-only and standard-plus-shifted processing produced identical allele-level calls and identical VAF estimates, including at the designated boundary positions.

This null result should not be interpreted as evidence that circular-aware processing is unnecessary. It shows that the particular benefit being tested, incremental boundary-specific variant recovery, was not measurable once downstream processing was held constant in this fixture. Read length, fragment length, variant position, aligner behavior, and the structure of the simulated reads were fixed in the experiment rather than varied independently. Different combinations of these factors may produce different effects at the artificial breakpoint.

The result is nevertheless useful because circular handling is often adopted for sound biological reasons without separately measuring the contribution of the shifted branch to final variant recovery. In MitoArc, the standard-plus-shifted procedure remains part of the production workflow because it provides complete canonical coverage without relying on the linear breakpoint. The benchmark result simply places a limit on the claim that can be made from the current data: shifted-coordinate processing was implemented and tested, but an additional boundary-specific recovery advantage was not demonstrated under the evaluated conditions.

### 4.3 Increasing sequencing depth improves some measurements but does not define a universal optimum

Sequencing depth affected both variant recovery and VAF estimation. GATK recall increased from 0.741 at 100× to 0.917 at 5000×, while its mean absolute VAF error decreased from 0.00884 to 0.00054. mutserve2 recall increased over the lower depth range but changed little beyond approximately 500× under the frozen configuration, whereas its VAF error continued to decrease. These results separate two effects that can otherwise be conflated: additional sequencing may improve quantitative allele-fraction estimation even when the number of recovered variants changes relatively little.

Other mtDNA studies have also treated depth as a methodological variable rather than assuming that all available mitochondrial reads should always be retained. MitoHPC down-samples mitochondrial reads to approximately 2000× before variant analysis to reduce computational cost while retaining the coverage required for its targeted heteroplasmy range (Battle et al. 2022). mtDNA-Server 2 similarly provides optional coverage subsampling and evaluated caller behavior at 100× and 2000× (Weissensteiner et al. 2024). A separate coverage study using mutserve2 found that reliability of short-read heteroplasmy detection depended jointly on VAF and sequencing depth, with low-frequency calls becoming more reliable as coverage increased (Slapnik et al. 2024).

These studies should not be used to define a common depth threshold because their read generation, truth sets, callers, filtering criteria, and heteroplasmy ranges differ. The MitoArc benchmark also deliberately included nominal VAFs as low as 0.5%, used achieved rather than requested VAF for quantitative evaluation, and introduced no background sequencing-error model. Its depth curves therefore describe the controlled experiment rather than a clinical limit of detection.

Computational cost also changed differently across resource dimensions. Between 100× and 5000×, mean elapsed time increased 3.56-fold, whereas peak working-directory storage increased approximately 20-fold; peak resident memory reached 1.4 GB. The strongest resource penalty at high depth was therefore temporary storage rather than a proportional increase in all computational measures. This observation supports exposing depth and resource use to the user rather than treating extremely high mitochondrial coverage as computationally neutral.

The present data are not sufficient to define an automatic downsampling threshold. The appropriate depth depends on the VAF range and variant classes of interest, the sequencing protocol, and the caller configuration. A future depth-aware mode would therefore require additional validation before it could discard reads on the basis of an assumed performance plateau.

### 4.4 Caller differences reflect both algorithmic behavior and configuration

GATK Mutect2 and mutserve2 produced different results when evaluated against the same truth panel. GATK recovered substantially more truth observations, while overall precision was similar between the callers. mutserve2 had a lower mean absolute VAF error among the alleles that it recovered, but this value was calculated over a smaller and different subset of the truth panel. These results therefore do not support reducing caller comparison to a single ranking.

Variant class was an important part of the difference. GATK recovered 172 of 216 single-base insertion truth observations, whereas none of the truth insertions were emitted by mutserve2 in the evaluated configuration. Diagnostic tracing showed that the insertions were already absent from the native mutserve2 output, rather than being removed during MitoArc normalization. The frozen v1.0.0 command did not enable mutserve2’s beta insertion and deletion options. This distinction is important because it makes the result configuration-specific rather than a statement about the intrinsic ability of mutserve2 to analyze indels.

The design of mtDNA-Server 2 illustrates the same broader point. Its analysis modes use mutserve2 for SNVs, Mutect2 for SNVs and indels, or a fusion mode in which mutserve2 SNVs are combined with Mutect2 indels (Weissensteiner et al. 2024). MitoHPC has also used both Mutect2 and mutserve, but with a different set of caller options and post-processing choices, including explicit mutserve insertion and deletion flags in the published implementation (Battle et al. 2022). Numerical performance estimates from different pipelines therefore cannot be interpreted independently of the exact caller mode in which they were generated.

MitoArc retains the two caller outputs separately rather than constructing a preferred or fused callset. The HG002 analysis provides a practical example of why this provenance is useful. GATK produced 35 variant calls and mutserve2 produced 21, with 16 shared sites, yet both callsets assigned H5a7 as the highest-ranked haplogroup. Their default consensus sequences were also sequence-identical, although the mutserve2 consensus retained a failed QC status because of a multiallelic conflict. A shared downstream summary can therefore coexist with meaningful differences in the evidence from which that summary was derived.

### 4.5 MitoArc complements existing mtDNA workflows by making analytical choices directly testable

MitoArc enters a field with several established human mitochondrial analysis pipelines. MToolBox introduced integrated mapping, NumtS filtering, heteroplasmy analysis, haplogroup prediction, and annotation; MitoHPC added high-throughput WGS analysis with mtDNA copy-number estimation, circular remapping, NUMT-read recovery, and sample-specific mitochondrial references; mtDNA-Server 2 provides scalable containerized analysis with mutserve2, Mutect2, haplogroup and contamination analysis, and interactive reporting; and MitoGEx provides an integrated graphical workflow covering preprocessing, GATK mitochondrial calling, annotation, haplogroup classification, and visualization (Battle et al. 2022; Calabrese et al. 2014; Jeenkeawpiam et al. 2026; Weissensteiner et al. 2024).

The distinguishing emphasis of the present study is therefore not the existence of an integrated mtDNA pipeline. MitoArc places several consequential analytical choices within one frozen workflow and evaluates them explicitly: mapping-context filtering is compared using an observability-aware NUMT challenge; standard-only and standard-plus-shifted processing are tested with matched downstream treatment; sequencing depth is evaluated against both variant metrics and computational cost; and the two caller outputs are preserved and assessed against the same allele-level truth. The null circular-coordinate result and the sensitivity loss associated with Strategy B are retained alongside positive findings rather than removed through post-benchmark retuning.

The HG002 experiment provides a complementary type of validation. The reduced alignment passed through the downstream workflow and produced coverage, caller-specific variants, NUMT evidence, haplogroup assignments, annotation, consensus sequences, plots, and reporting. However, it contained only selected nuclear regions and no independent mtDNA truth set. It therefore supports real-data operation of the workflow rather than accuracy or complete-WGS NUMT performance. Maintaining this distinction prevents technical workflow operation from being conflated with truth-based accuracy.

Reproducibility in this setting also concerns more than software installation. Reference sequence, alignment context, caller parameters, VAF thresholds, normalization rules, and treatment of ambiguous evidence can all alter the set of reported mitochondrial variants. MitoArc records these choices while retaining machine-readable caller and benchmark outputs, rather than reducing them to a final report alone. This is particularly important when different analytical routes can converge on the same downstream haplogroup or consensus sequence despite differing at the underlying variant level.

### 4.6 Limitations and future evaluation

Several limitations define the scope of the present study. The controlled benchmark used synthetic paired-end reads generated from fixed mitochondrial truth positions and three stochastic technical replicates. The primary panel contained SNVs and single-base insertions but no deletion truth alleles, and no stochastic background sequencing-error model was added. The synthetic nuclear background included short neutral autosomal scaffolds and designed mitochondrial-homologous contigs rather than the sequence complexity of a complete human genome. The benchmark therefore isolates selected sources of analytical behavior but does not reproduce the full range of errors, repeats, structural variation, and polymorphic NUMTs encountered in human WGS.

The NUMT conclusions are similarly bounded. Strategy B was tested against engineered NUMT and ambiguous challenges in a reference-defined setting, while Strategy C currently performs fragment-level support classification only for SNVs. The benchmark does not establish universal NUMT suppression, and recent pangenome evidence makes clear that reference and non-reference NUMTs constitute a broader problem than can be represented by the current synthetic model (Fu et al. 2026).

Caller conclusions apply to the frozen v1.0.0 configurations. In particular, mutserve2 insertion and deletion modes were disabled, so the absence of primary-panel insertion calls cannot be generalized to other mutserve2 modes. Likewise, the depth curves reflect these caller settings, the benchmark’s allele spectrum, and its lack of background sequencing error. The three replicates are technical rather than biological replicates, and the study was intentionally analyzed descriptively without inferential hypothesis testing.

The HG002 real-data validation is also limited by its reduced nuclear context. HG002 retained complete chrM data but only selected regions from chromosomes 1, 2, and 17, preventing complete autosomal coverage estimation and complete-WGS NUMT assessment. The study also did not perform a publication-scale complete-human FASTQ run, clinical validation, long-read validation, or testing across a large set of biological samples and haplogroups. Resource measurements were obtained from the controlled executions and should not be interpreted as universal hardware requirements.

Future evaluation should therefore extend the workflow to complete human WGS inputs, more diverse mitochondrial sequence backgrounds, additional read lengths and fragment distributions, broader indel classes, and more realistic sequencing-error models. Long-read or pangenome-informed NUMT resources would also provide a stronger basis for testing nuclear-mitochondrial origin than a single reference genome. These extensions would allow the current methodological observations to be tested under a wider range of biological and sequencing conditions without assuming that the effects measured here generalize beyond the present benchmark.

## 5. Conclusion

MitoArc provides a reproducible workflow for human short-read mitochondrial genome analysis from paired-end FASTQ, BAM, and CRAM inputs while preserving caller-specific evidence and the analytical context in which mitochondrial variants are produced. For FASTQ input, the workflow performs full-reference mapping before mitochondrial read extraction, while pre-aligned BAM/CRAM inputs retain their available mapping context. Downstream analysis combines circular-coordinate processing, independent GATK Mutect2 and mutserve2 calling, NUMT evidence assessment, haplogroup and contamination analysis, functional annotation, consensus generation, and portable reporting within a version-controlled Nextflow implementation.

Controlled benchmarking showed that the analytical choices surrounding mtDNA variant detection can have measurable and sometimes opposing effects. Requiring confident mitochondrial mapping removed the observable synthetic NUMT-associated false-positive signals in the evaluated benchmark but reduced recovery of true mitochondrial variants. Increasing sequencing depth improved GATK recall and reduced allele-fraction error, while computational requirements, particularly temporary storage, continued to rise. GATK and mutserve2 showed different recovery patterns under the same truth framework, and the absence of mutserve2 insertion calls reflected the configuration tested. In contrast, matched GATK standard-only and standard-plus-shifted processing produced identical variant recovery and VAF estimates in the evaluated circular-boundary fixture, providing no evidence for an incremental boundary-specific gain under those conditions.

These findings support an approach to mtDNA analysis in which mapping context, caller configuration, sequencing depth, and reference handling are recorded and evaluated rather than hidden behind a single final callset. MitoArc is not intended to define a universally optimal caller, NUMT filter, or sequencing depth. Instead, it provides a reproducible basis for analyzing human short-read mtDNA while retaining the evidence and methodological choices needed to interpret the resulting variants.

## Supporting information

Suppelementary Info

## Supplementary Information

Online Resource 1 contains supplementary methods, figures, tables, and supporting analyses for the MitoArc workflow and benchmark.

## Author contributions

Sarvesh Galgale contributed to the methodology, software development, formal analysis, validation, investigation, preparation of the original draft, and review and editing of the manuscript. Isha Vats contributed to the methodology, software development, formal analysis, validation, investigation, preparation of the original draft, and review and editing of the manuscript. Agnik Haldar contributed to the conceptualization, methodology, supervision, and review and editing of the manuscript. All authors read and approved the final manuscript.

## Data availability

MitoArc v1.0.0 is available from GitHub at https://github.com/sarvezh1/mitoarc/tree/v1.0.0 and is archived on Zenodo at https://doi.org/10.5281/zenodo.23154158. The archived release contains the workflow source code, configuration files, supporting scripts, validation resources, and machine-readable benchmark data used in this study. Data underlying selected results reported in the main manuscript and Online Resource 1 are provided in the paper_data/ directory at https://github.com/sarvezh1/mitoarc/tree/v1.0.0/paper_data. The public HG002/NA24385 Illumina HiSeq 300× GRCh38 alignment used for regional retrieval is available through the Genome in a Bottle alignment index at https://github.com/genome-in-a-bottle/giab_data_indexes/blob/master/AshkenazimTrio/alignment.index.AJtrio_Illumina300X_wgs_novoalign_GRCh37_GRCh38_NHGRI_07282015.

## Statements and Declarations

### Ethics approval and consent to participate

Not applicable. This computational study used synthetic benchmark data and publicly available, de-identified sequencing data from the GIAB HG002 reference sample. No participants were recruited, and no identifiable private information was used.

### Consent for publication

Not applicable.

