## Supplementary material for "MitoArc: A Reproducible Short-Read Workflow for Human Mitochondrial Genome Analysis with Circular-Coordinate and NUMT-Aware Evaluation": Suppelementary Info

**Journal:** Network Modeling Analysis in Health Informatics and Bioinformatics

Sarvesh Galgale¹,², Isha Vats³, Agnik Haldar¹,²

¹Department of Biotechnology, School of Applied Sciences, REVA University, Kattigenahalli, Yelahanka, Bengaluru, Karnataka 560064, India

²Center for Biotechnology and Molecular Biology, REVA Research Centre, REVA University, Kattigenahalli, Yelahanka, Bengaluru, Karnataka 560064, India

³Kusuma School of Biological Sciences, Indian Institute of Technology Delhi, Hauz Khas, New Delhi 110016, India

Sarvesh Galgale and Isha Vats contributed equally to this work.

**Corresponding author:** Agnik Haldar

### **Supplementary Methods**

#### **S1. Controlled benchmark design and technical replicates**

The controlled short-read mitochondrial benchmark contained 36 mitochondrial truth alleles in each dataset: 24 SNVs and 12 single-base insertions. The primary truth panel contained no deletion alleles. Six nominal variant allele fraction (VAF) strata were evaluated: 0.005, 0.015, 0.035, 0.075, 0.25, and 0.75. Each VAF stratum included six allele archetypes spanning the designed variant and mapping contexts. Three independently seeded technical replicates were generated at target mitochondrial depths of 100×, 250×, 500×, 1000×, 2000×, and 5000×. These replicates capture stochastic technical variation at fixed truth positions and are not biological replicates. This design produced 18 physical workflow executions and 648 mitochondrial truth observations. Nominal VAF defined the experimental design strata. Quantitative VAF-error calculations used the achieved VAF in each generated dataset, and error was evaluated among recovered truth observations. Benchmark summaries are descriptive; no inferential tests were introduced, and the tested depth range was not used to define an optimal sequencing depth.

#### **S2. NUMT challenge observability and mapping-context evaluation**

The benchmark included designed NUMT-like and ambiguous-context challenge signals for false-positive assessment. Each physical challenge was tracked through five stages: designed, generated in the synthetic reads, mapped during full-reference alignment, extracted into the mitochondrial analysis path, and observable for downstream NUMT-associated false-positive assessment. Counts were assigned to physical challenge identities rather than caller-specific copies. All 216 designed challenges were generated and mapped; 198 were extracted and observable. The 18 challenges that did not enter the mitochondrial analysis path were excluded from downstream false-positive opportunities and were not counted as filtering successes. Strategy A represents the baseline caller output without an additional mapping-context filter. Strategy B retains read names supported by a qualifying primary mitochondrial alignment before repeating the mitochondrial calling paths. False positives in this comparison use OBSERVABLE_NUMT_ASSOCIATED_FP, not the overall caller metric TOTAL_EXACT_TRUTH_FP. Mapping-context filtering eliminated the observable synthetic NUMT-associated false positives in this benchmark, but reduced recall. This controlled result does not establish a universal benefit and is not an estimate of natural NUMT burden.

#### **S3. Strategy C evidence classification and unresolved non-SNV support**

Strategy C is an evidence-classification view rather than an independent variant caller. For eligible SNV support, it links supporting fragment names to retained mapping records and classifies the available mapping evidence as LOW, AMBIGUOUS, or HIGH. At the variant level, an SNV with at least two supporting fragments and no LOW evidence is assigned FLAGGED_NUMT_EVIDENCE; other called SNVs are assigned RETAINED. Strategy C does not remove records from caller VCFs. Fragment-level allele support is restricted to SNVs in MitoArc v1.0.0. Non-SNV records are outside this classification scope and may receive UNRESOLVED_SUPPORT. The six retained diagnostic non-SNV traces shown in Figure S2 each contain nonzero ALT-supporting fragments in the mitochondrial alignment despite a final UNRESOLVED_SUPPORT status. These traces show that this status can reflect classifier scope rather than absence of supporting reads, but the observation is limited to the six traced examples. The complete read-level Strategy C evidence table from the original benchmark execution was not retained. Accordingly, this supplementary file reports the implemented classification logic, retained class summaries, and six available diagnostic traces rather than reconstructing unavailable read-level evidence.

#### **S4. Matched GATK circular-coordinate evaluation**

The matched GATK circular-coordinate experiment compared standard-only processing with standard-plus-shifted processing while holding downstream calling, filtering, and normalization constant. It included three technical replicates at mitochondrial depths of 250×, 1000×, and 5000× and nominal VAFs of 0.015, 0.075, and 0.25. The comparison comprised 108 allele-level truth observations representing boundary and non-boundary SNVs and single-base insertions. Calls were evaluated by exact normalized allele matching. Recovery was compared for every truth observation, and caller-reported VAFs were compared within recovered allele pairs. All 108 allele-level calling outcomes were identical between processing modes. Recovered-allele called VAF values were also identical between modes. Pooled recall differences were zero for boundary SNVs, boundary insertions, non-boundary SNVs, and non-boundary insertions. No incremental boundary-specific recovery was observed under the matched conditions tested. This experiment isolates the incremental effect under the specified matched conditions and does not establish that circular-coordinate handling is unnecessary.

#### **S5. Computational resource measurements and software versions**

Computational resources were measured for complete physical workflow executions rather than for individual caller branches. Three technical replicate executions were summarized at each target depth. Measurements included elapsed wall time, CPU hours, peak resident set size (RSS), peak work-directory storage, and final output size. Elapsed time, CPU hours, work-directory size, and output size are reported as means across the three executions at each depth; peak RSS is the maximum observed across replicates. Each execution included both caller outputs and the NUMT comparison branches, so the measurements are not caller-specific. Decimal megabytes are used for displayed storage values, while the machine-readable resource table retains precise byte values. The results describe the measured benchmark executions and are not used to recommend an optimal sequencing depth or to assert a linear scaling model. Table S1 lists the verified software versions, including MitoArc v1.0.0. Table S5 reports the depth-specific resource measurements.

### **Supplementary Figures**

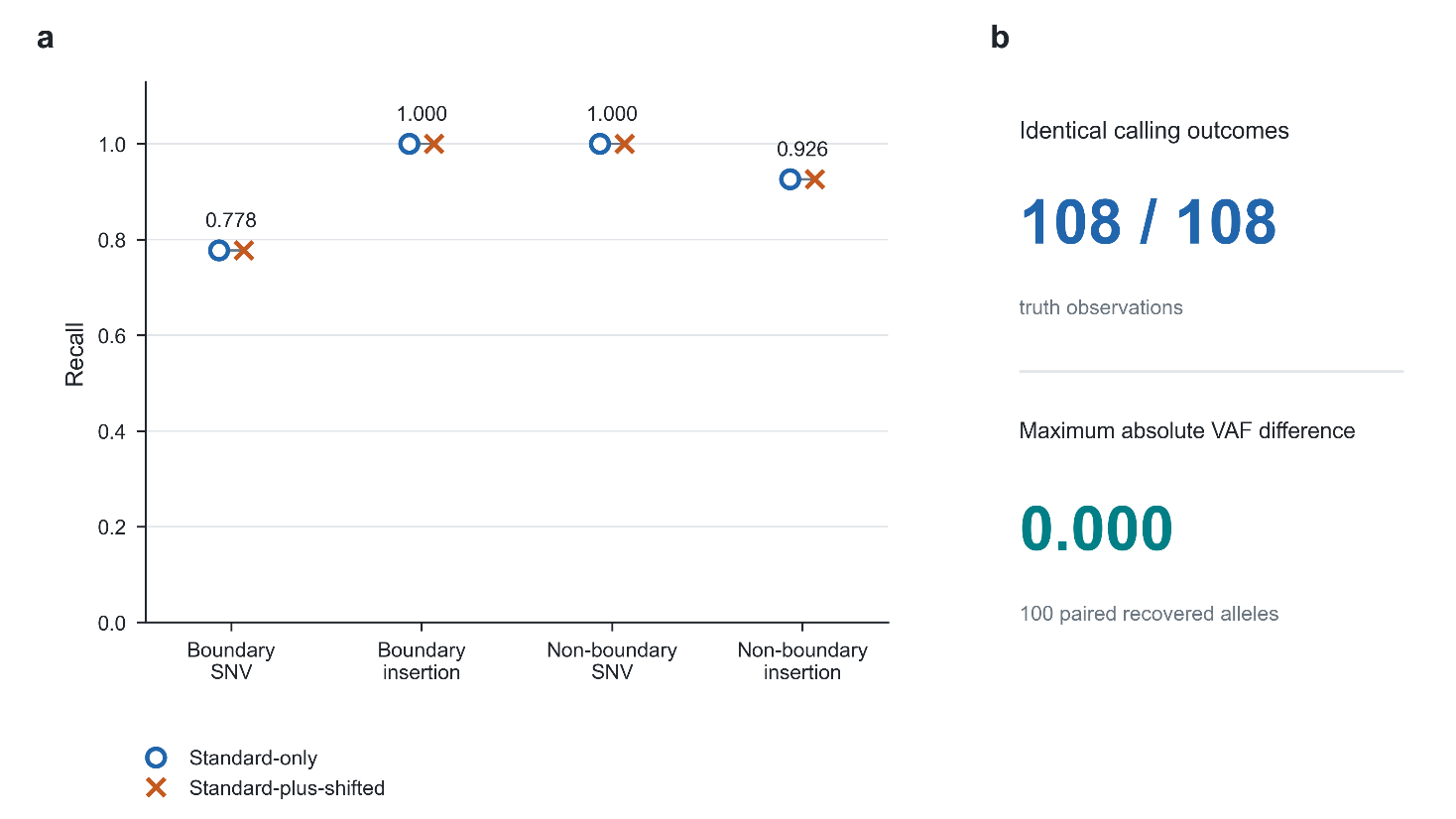

**Fig. S1** Matched GATK circular-coordinate evaluation. (a) Recall for boundary and non-boundary SNVs and insertions under matched standard-only and standard-plus-shifted processing. Identical values are shown for both modes. (b) Calling outcomes were identical for 108/108 allele-level observations, and the maximum absolute called-VAF difference among recovered allele pairs was zero (max |ΔVAF| = 0.000). No incremental boundary-specific recovery was observed under the matched conditions tested

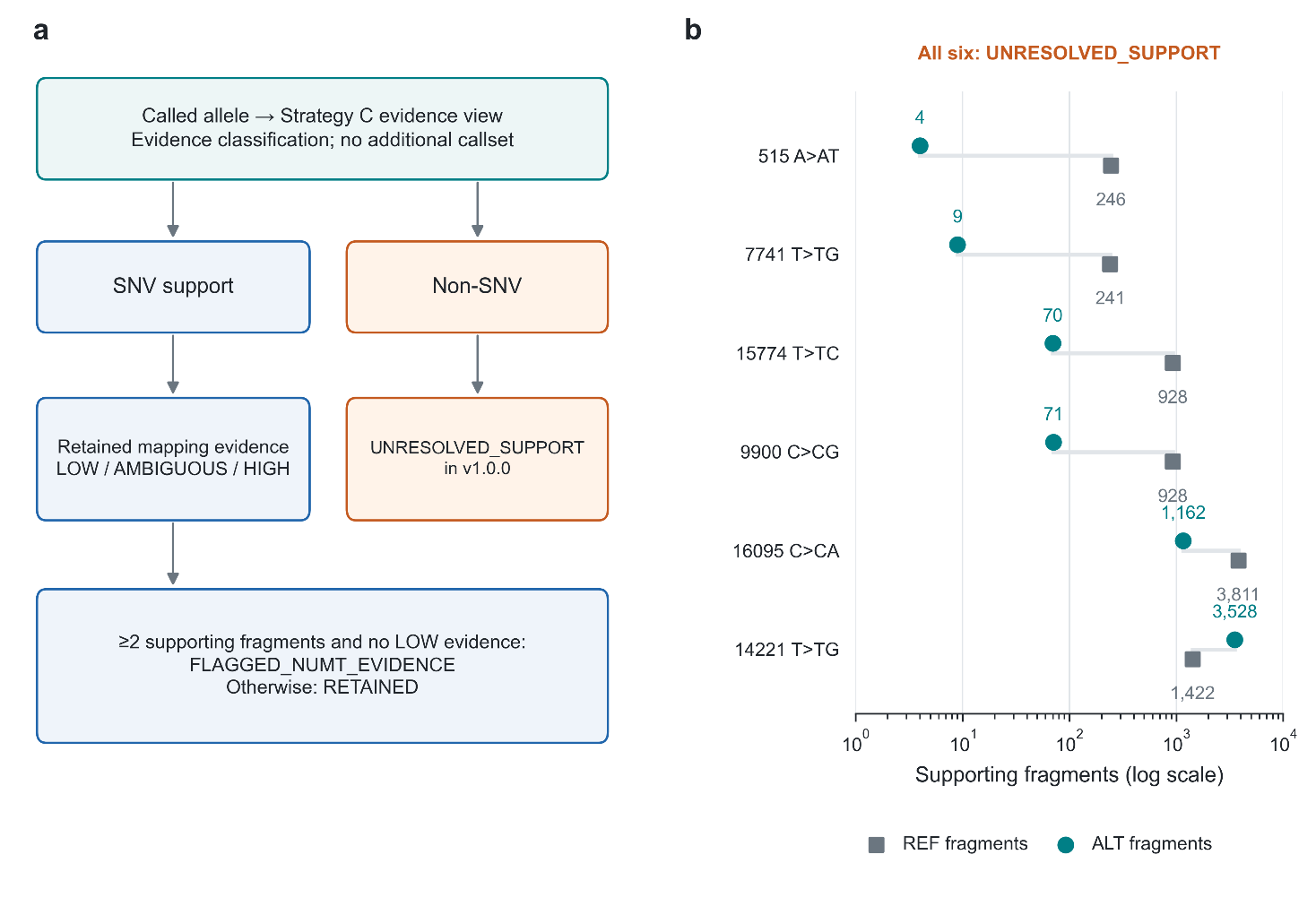

**Fig. S2** Strategy C evidence scope and unresolved non-SNV support. (a) Strategy C classifies retained mapping evidence for eligible SNV support and is an evidence view rather than an independent caller. Non-SNV records are outside the v1.0.0 fragment-level classification scope. (b) The six retained diagnostic non-SNV traces show REF- and ALT-supporting fragments in the mitochondrial alignment and a final UNRESOLVED_SUPPORT status. These examples show that UNRESOLVED_SUPPORT can reflect classifier scope rather than absence of supporting reads; this interpretation is limited to the six retained traces

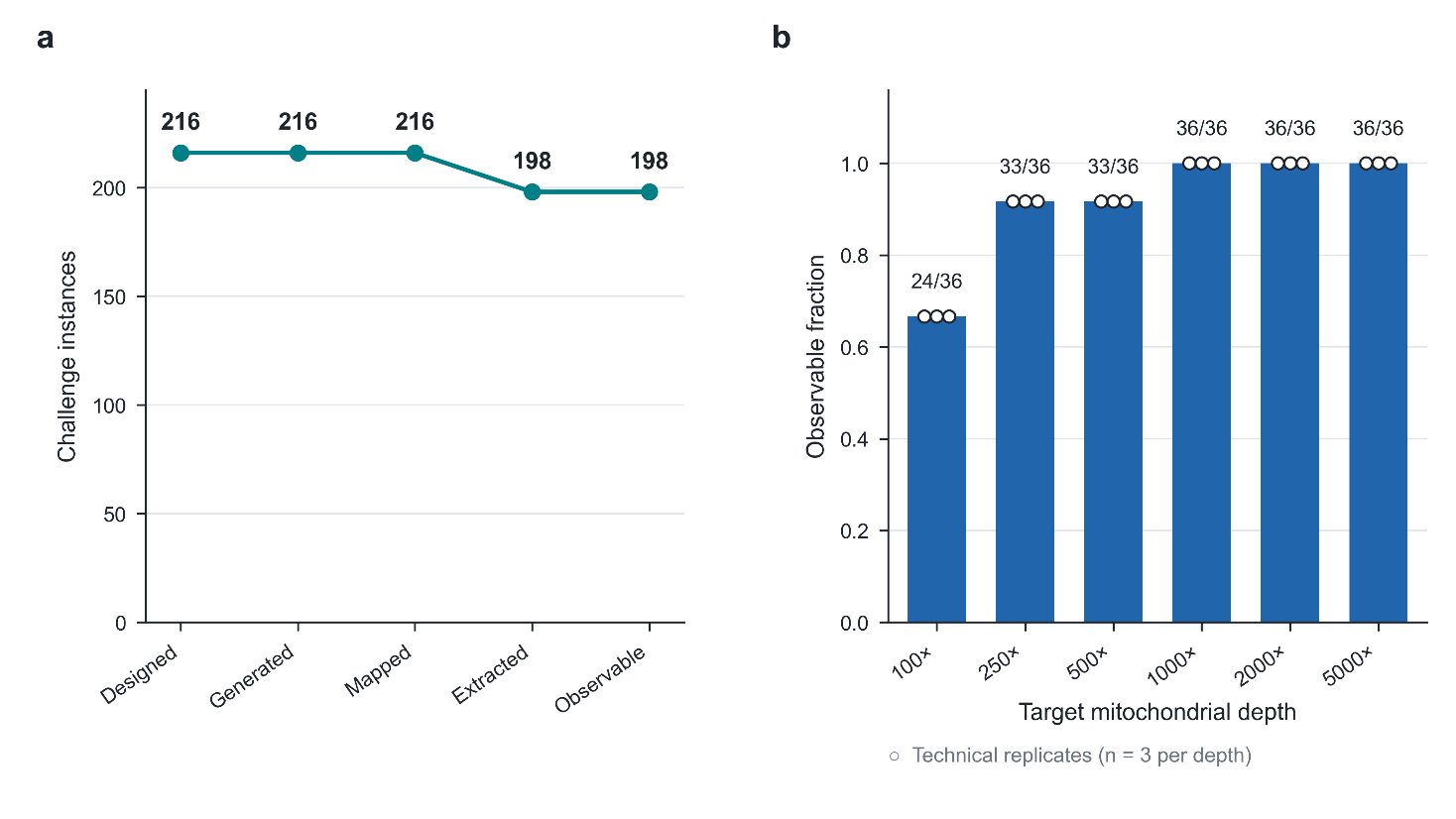

**Fig. S3** NUMT challenge observability. (a) Progression of physical challenge instances from designed through generated, mapped, extracted, and observable stages. (b) Observable fraction by target mitochondrial depth; points show the three technical replicates. Eighteen of 216 designed challenge instances did not remain observable after mitochondrial extraction and were not counted as downstream-filtering successes

### **Supplementary Tables**

**Table S1** Software and version manifest.

| **Software** | **Version** |
| --- | --- |
| MitoArc | 1.0.0 |
| Nextflow | 26.04.6 |
| GATK | 4.6.2.0 |
| mutserve2 | 2.0.3 |
| HaploGrep | 3.3.2 |
| Haplocheck | 1.3.3 |
| MultiQC | 1.27.1 |
| matplotlib | 3.10.5 |
| BWA-MEM2 | 2.2.1 |
| samtools | 1.20 |
| bcftools | 1.20 |
| mosdepth | 0.3.14 |
| fastp | 0.23.4 |
| FastQC | 0.12.1 |

**Table S2** Caller performance and VAF error by sequencing depth.

| **Caller** | **Depth** | **TP** | **FP** | **FN** | **Precision** | **Recall** | **F1** | **Mean absolute VAF error** |
| --- | --- | --- | --- | --- | --- | --- | --- | --- |
| GATK Mutect2 | 100× | 80 | 12 | 28 | 0.869565 | 0.740741 | 0.800000 | 0.008836 |
| GATK Mutect2 | 250× | 87 | 12 | 21 | 0.878788 | 0.805556 | 0.840580 | 0.003417 |
| GATK Mutect2 | 500× | 91 | 6 | 17 | 0.938144 | 0.842593 | 0.887805 | 0.002199 |
| GATK Mutect2 | 1000× | 95 | 3 | 13 | 0.969388 | 0.879630 | 0.922330 | 0.001414 |
| GATK Mutect2 | 2000× | 96 | 0 | 12 | 1.000000 | 0.888889 | 0.941176 | 0.000740 |
| GATK Mutect2 | 5000× | 99 | 0 | 9 | 1.000000 | 0.916667 | 0.956522 | 0.000544 |
| mutserve2 | 100× | 36 | 0 | 72 | 1.000000 | 0.333333 | 0.500000 | 0.004772 |
| mutserve2 | 250× | 50 | 3 | 58 | 0.943396 | 0.462963 | 0.621118 | 0.001981 |
| mutserve2 | 500× | 56 | 5 | 52 | 0.918033 | 0.518519 | 0.662722 | 0.001558 |
| mutserve2 | 1000× | 57 | 3 | 51 | 0.950000 | 0.527778 | 0.678571 | 0.001313 |
| mutserve2 | 2000× | 55 | 3 | 53 | 0.948276 | 0.509259 | 0.662651 | 0.000781 |
| mutserve2 | 5000× | 57 | 3 | 51 | 0.950000 | 0.527778 | 0.678571 | 0.000649 |

Note. TP, FP, and FN are pooled across the three technical replicates at each depth; precision, recall, and F1 are calculated from these pooled counts. FP uses TOTAL_EXACT_TRUTH_FP. Mean absolute VAF error is based on achieved VAF among recovered truth observations and summarizes replicate-level errors; recovered truth subsets differ between callers. mutserve2 results apply to the frozen MitoArc v1.0.0 configuration, in which beta insertion and deletion modes were not enabled.

**Table S3** NUMT Strategy A/B results and observability.

| **Replicate** | **Designed NUMT/ambiguous challenges** | **Observable challenges** | **Strategy A TP** | **Strategy A observable NUMT-associated FP** | **Strategy A FN** | **Strategy A Recall** | **Strategy B TP** | **Strategy B observable NUMT-associated FP** | **Strategy B FN** | **Strategy B Recall** |
| --- | --- | --- | --- | --- | --- | --- | --- | --- | --- | --- |
| R1 | 72 | 66 | 65 | 11 | 7 | 0.902778 | 58 | 0 | 14 | 0.805556 |
| R2 | 72 | 66 | 64 | 11 | 8 | 0.888889 | 57 | 0 | 15 | 0.791667 |
| R3 | 72 | 66 | 64 | 11 | 8 | 0.888889 | 57 | 0 | 15 | 0.791667 |
| Pooled | 216 | 198 | 193 | 33 | 23 | 0.893519 | 172 | 0 | 44 | 0.796296 |

Note. Each technical replicate contained 72 true mitochondrial observations used for TP/FN recall assessment and 72 designed NUMT/ambiguous-context challenges used for false-positive assessment; 66 of the latter were observable after mitochondrial extraction. False positives use OBSERVABLE_NUMT_ASSOCIATED_FP. Mapping-context filtering eliminated the observable synthetic NUMT-associated false positives in this benchmark, but reduced recall. In the pooled row, the corresponding totals are 216 true mitochondrial observations, 216 designed NUMT/ambiguous-context challenges, and 198 observable challenges.

**Table S4** Matched GATK circular-coordinate results.

| **Boundary class** | **Variant class** | **Truth observations** | **Standard-only TP** | **Standard-only FN** | **Standard-only Recall** | **Standard-plus-shifted TP** | **Standard-plus-shifted FN** | **Standard-plus-shifted Recall** | **Recall delta** | **Maximum absolute VAF difference** |
| --- | --- | --- | --- | --- | --- | --- | --- | --- | --- | --- |
| Boundary | SNV | 27 | 21 | 6 | 0.777778 | 21 | 6 | 0.777778 | 0.000 | 0.000 |
| Boundary | Insertion | 27 | 27 | 0 | 1.000000 | 27 | 0 | 1.000000 | 0.000 | 0.000 |
| Non-boundary | SNV | 27 | 27 | 0 | 1.000000 | 27 | 0 | 1.000000 | 0.000 | 0.000 |
| Non-boundary | Insertion | 27 | 25 | 2 | 0.925926 | 25 | 2 | 0.925926 | 0.000 | 0.000 |

Note. Calling, filtering, and normalization were matched between processing modes. Recall delta is the standard-plus-shifted recall minus the standard-only recall. Maximum absolute VAF difference is the largest absolute difference between paired caller-reported VAFs among truth alleles recovered in both modes.

**Table S5** Computational resource measurements.

| **Depth** | **Mean elapsed time, s** | **Mean CPU hours** | **Maximum peak RSS, MB** | **Mean peak work-directory size, MB** | **Mean final output size, MB** |
| --- | --- | --- | --- | --- | --- |
| 100× | 332.6 | 0.102 | 1100 | 58.5 | 18.7 |
| 250× | 357.1 | 0.109 | 1100 | 83.4 | 21.9 |
| 500× | 370.3 | 0.116 | 1100 | 141.1 | 26.6 |
| 1000× | 458.7 | 0.131 | 1100 | 256.3 | 34.9 |
| 2000× | 591.1 | 0.149 | 1200 | 485.0 | 50.7 |
| 5000× | 1184.3 | 0.237 | 1400 | 1170.8 | 96.4 |

Note. Values summarize three technical replicate workflow executions at each target depth. Elapsed time, CPU hours, work-directory size, and output size are means; peak RSS is the maximum observed across replicates. Measurements represent complete physical workflow executions, including both caller outputs and NUMT comparison branches, and are not caller-specific. Decimal megabytes are reported.

### **Supplementary Data Files**

Machine-readable values underlying selected benchmark results reported in the manuscript and this Online Resource are available in the paper_data/ directory of MitoArc v1.0.0:

<https://github.com/sarvezh1/mitoarc/tree/v1.0.0/paper_data>

The same files are preserved in the archived release at:

<https://doi.org/10.5281/zenodo.23154158>

- software_versions.tsv: verified software and version manifest.
- caller_depth_metrics.tsv: caller TP, FP, FN, precision, recall, F1, and achieved-VAF error by sequencing depth. False positives use TOTAL_EXACT_TRUTH_FP.
- numt_observability.tsv: designed, generated, mapped, extracted, and observable NUMT challenge counts by depth and replicate.
- numt_strategy_metrics.tsv: Strategy A/B TP, observable NUMT-associated FP, FN, and recall for technical replicates and pooled results. False positives use OBSERVABLE_NUMT_ASSOCIATED_FP.
- circular_matched_metrics.tsv: matched standard-only and standard-plus-shifted circular-coordinate results.
- resource_metrics.tsv: elapsed time, CPU hours, peak RSS, work-directory size, and output size for complete physical workflow executions.
